# Inhibition of KDM5A/B promotes antitumor innate immune responses in HHV-8/KSHV-positive B-cell lymphomas

**DOI:** 10.64898/2026.01.28.702275

**Authors:** Dawei Zhou, Guillaume N Fiches, Zhenyu Wu, Suha Eleya, Youngmin Park, Jinshan He, Kateepe ASN Shanaka, Thurbu T Lepcha, Yan Liu, Jason Oliva, Kathryn Lurain, Jae U Jung, Jun Qi, Weiqiang J Zhao, Jian Zhu, Netty G Santoso

## Abstract

Histone methylation is a dynamic and reversible epigenetic modification that critically controls the progression of human diseases, including infections and cancers. Here we reported that histone lysine demethylases (KDMs) in the KDM5 family KDM5A/B play profound roles in suppressing lytic reactivation of oncogenic human herpesvirus 8 (HHV-8), i.e., Kaposi’s sarcoma-associated herpesvirus (KSHV), as well as antiviral/antitumor innate immune responses in KSHV-infected B-cell lymphomas. We showed that KSHV lytic replication decreases KDM5A/B protein stability by enhancing their K-48 linked polyubiquitination while KDM5A/B depletion facilitates KSHV lytic reactivation. Mechanistic studies illustrated that KDM5A/B associate with KSHV LANA protein and dampen its chromatin association at both KSHV viral lytic promoter and promoters of antitumor immune-responsive genes (IRGs). In comparisons to normal B cells, KDM5A/B expression significantly increased in B-cell lymphoma cells, including KSHV-positive primary effusion lymphoma (PEL). We demonstrated that KDM5A/B inhibition remarkably induces both KSHV lytic reactivation and innate immune responses in PEL cells, resulting in a strong viral oncolytic effect, both *in vitro* in cell cultures and *in vivo* using a PEL xenograft mouse model. Overall, our studies identified the novel functions of KDM5A/B to silence KSHV lytic replication and antiviral/antitumor innate immune responses, which can be blocked to benefit the treatment of KSHV-associated B-cell lymphomas that are usually aggressive and difficult to treat.

## Introduction

HHV-8/KSHV is a human gamma-herpesvirus and an etiological agent of multiple malignancies, including Kaposi’s sarcoma(1), primary effusion lymphoma (PEL)(2), and multicentric Castleman’s disease (CAD)(3). Once KSHV enters the host cells, it goes through brief lytic replication and establishes latent infection by maintaining viral genomes as extra-chromosomal circular episomes in nuclei with most viral lytic genes silenced(4,5). KSHV lytic reactivation can be triggered by certain stimuli so that new progeny viruses are generated for viral propagation(6). Such latent/lytic switch of KSHV is delicately controlled by both viral and host regulators, including KSHV-encoded latency-associated nuclear antigen (LANA) and replication and transcription activator (RTA) proteins, host transcription and epigenetic factors, and host factors that participate in immune responses(7). KSHV latent/persistent infection is known to promote cell survival, proliferation and immune evasion, eventually leading to tumor initiation and progression(8). On the other hand, it has been tempted to induce KSHV lytic reactivation in KSHV-positive tumor cells so that viral oncolysis effect is achieved(9). Therefore, it is critical to elucidate profound KSHV-host interactions, especially epigenetic regulatory mechanisms that govern viral latent/lytic switch and persistent infection, which will benefit the therapeutic development for treating KSHV-associated malignancies.

Earlier studies have revealed the functional relevance of epigenetic regulation, including histone methylation/demethylation modifications, in KSHV-host interactions(10–13). However, only limited host factors participating in these epigenetic processes have been characterized, which have primarily focused on those targeting histone methylation sites that associate with transcriptional inactivation, such as H3K9 and H3K27. On the contrary, it has been barely examined regarding the contributions of histone methylation sites that associate with transcriptional activation (H3K4, H3K36) as well as modification enzymes targeting these sites to KSHV viral infection and/or KSHV-induced tumorigenesis. The KDM5 family members (KDM5A-D) are capable of removing methyl groups from the tri- and di-methylation of lysine 4 of histone H3 (H3K4me3/me2) that usually peak at the promoter regions of actively transcribed genes, which generally results in gene suppression phenotype(14). In particular, KDM5A and KDM5B (KDM5A/B) are ubiquitously expressed and play a critical role in controlling cell proliferation and differentiation. There is strong evidence that KDM5A/B are oncogenes overexpressed, amplified, or mutated in multiple types of human cancers(15). Based on the continuous efforts of our earlier KDM5A/B studies(16–18), we demonstrated that KSHV lytic program targets KDM5A/B for proteasome degradation while KDM5A/B suppresses KSHV lytic reactivation and antiviral/antitumor innate immune responses in KSHV-positive B-cell lymphomas, which can be inhibited to induce viral oncolysis effect *in vitro* and *in vivo*.

## Results

### KSHV lytic program targets KDM5A/B for proteasome degradation

An early proteomic study has implicated that KSHV lytic replication may lead to the reduction of host proteins(17). We first evaluated whether KSHV impacts KDM5 proteins. We relied on the PEL cell line BCBL-1 harboring the latently infected KSHV genome while also carrying an integrated cassette for doxycycline (Dox) induced RTA expression (TREx.BCBL-1.RTA). Dox treatment potently reactivated latent KSHV in TREx.BCBL-1.RTA cells (**Fig S1A**), which indeed dramatically reduced the protein level of KDM5A/B (**Fig 1A**) and KDM5C (**Fig S1B**). In contrast, their mRNA levels were not significantly affected (**Fig 1B, S1C**). Dox treatment had no such effects on the BJAB cells, a KSHV-negative Burkitt lymphoma (BL) cell line (**Fig 1C, 1D, S1D, S1E**). We further determined KSHV’s impact on KDM5 proteins during its *de novo* infection. The telomerase-immortalized human endothelial (TIME) cells were spinoculated with KSHV BAC16 viruses (**Fig S1F, S1G**), which also dramatically decreased the protein level of KDM5A/B (**Fig 1E**) and KDM5C (**Fig S1H**) while only slightly reducing their mRNA level (**Fig 1F, S1I**).

**Figure 1.**
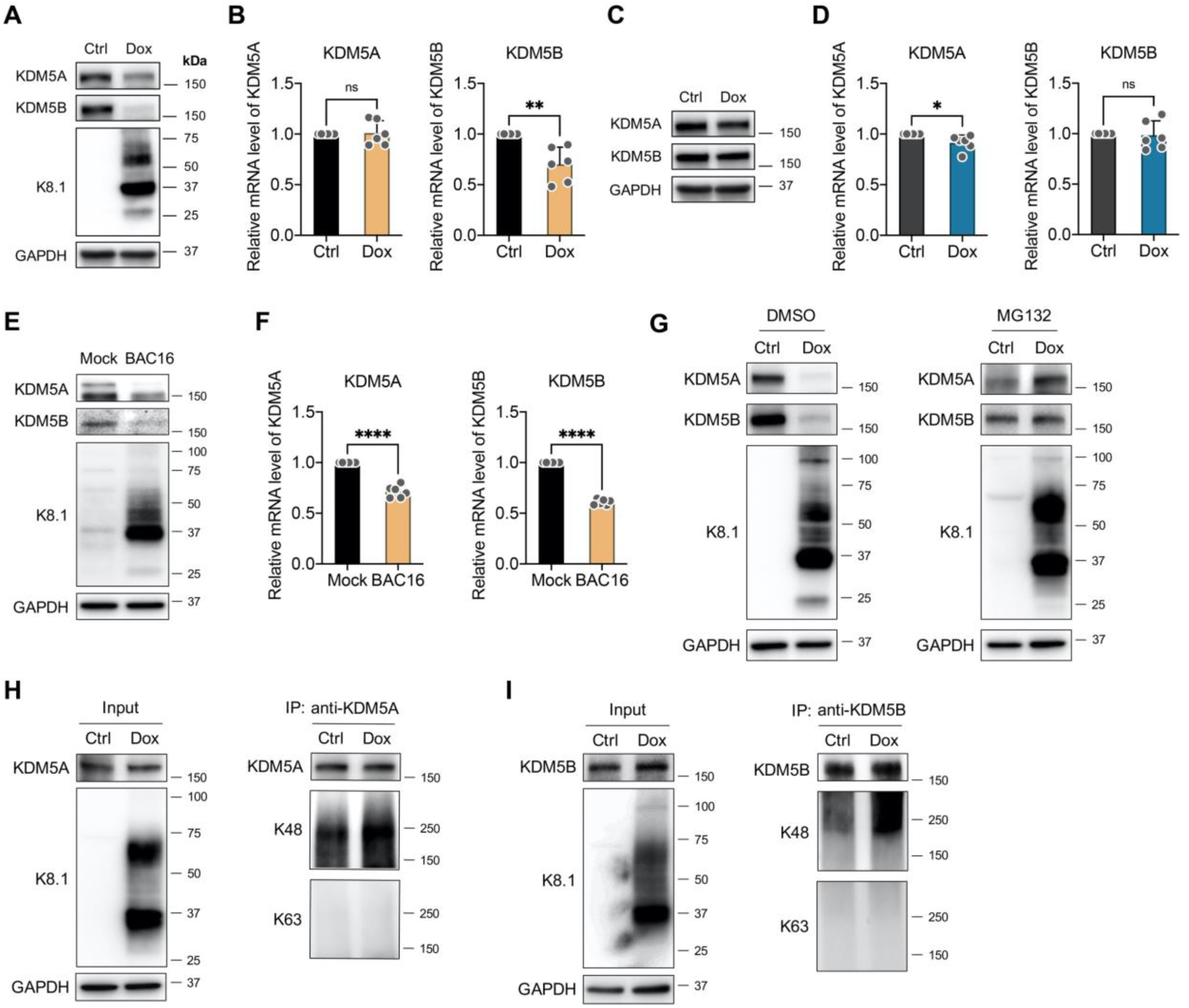
KSHV lytic program targets KDM5A/B for proteasome degradation. (A) TREx.BCBL-1.RTA cells were treated with doxycycline (Dox, 1 µg/mL) for 2 days, followed by protein immunoblotting to quantify KDM5A/B proteins and KSHV lytic K8.1 protein. (B) KDM5A/B mRNA level in cell samples (A) was measured by RT-qPCR. (C) BJAB cells were treated similarly as (A) and analyzed by protein immunoblotting to quantify KDM5A/B proteins. (D) KDM5A/B mRNA level in cell samples (C) was measured by RT-qPCR. (E) TIME cells were spinoculated with KSHV BAC16 viruses. At 2 days post of infection (dpi), cells were collected and analyzed by protein immunoblotting to quantify KDM5A/B proteins and KSHV lytic K8.1 protein. (F) KDM5A/B mRNA level in cell samples (E) was measured by RT-qPCR. (G) TREx.BCBL-1.RTA cells were treated with Dox in the absence or presence of MG132 (1 µM) for 2 days, followed by protein immunoblotting to quantify KDM5A/B proteins and KSHV lytic K8.1 protein. (H, I) TREx.BCBL-1.RTA cells were treated with Dox in the absence or presence of MG132 (1 µM) for 2 days, followed by protein immunoprecipitation (IP) of KDM5A (H) or KDM5B (I) using their specific antibodies, followed by protein immunoblotting to quantify K48 or K63-linked polyubiquitination of KDM5A/B. Results were calculated from three independent experiments and presented as Mean ± SD (ns: not significant, * *P* < 0.05, ** *P* < 0.01, **** *P* < 0.0001; unpaired, two-tailed Student’s *t*-test).

Based on these results, we hypothesized that KDM5 protein stability is affected by KSHV lytic program. To prove this, we treated the TREx.BCBL-1.RTA cells with a proteasome inhibitor MG132 (+/− Dox). MG132 restored the KDM5A/B protein level that was reduced due to Dox treatment (**Fig 1G**). Consistently, we showed that the K48-linked polyubiquitination of KDM5A/B is much higher in Dox-treated *vs* un-treated TREx.BCBL-1.RTA cells in the presence of MG132 (**Fig 1H, 1I**), while their K63-linked polyubiquitination was not detected. Early studies indicated that KSHV RTA promotes the protein degradation of certain host factors through upregulation of their K48-linked polyubiquitination(19,20). We employed the KSHV-negative BJAB cells carrying a cassette for Dox induced expression of 3 × FLAG-tagged RTA (TREx.BJAB.3FLAG.RTA) to determine whether RTA directly mediates KDM5 protein degradation. However, Dox treatment failed to decrease KDM5 proteins in these cells (**Fig S1J**). Transfection of RTA cDNA in HEK293T cells also failed to do so with or without MG132 (**Fig S1K**). Overall, our results demonstrated that KSHV lytic replication promotes proteosome degradation of KDM5 proteins (KDM5A-C), which may depend on other KSHV viral lytic components/events beyond RTA.

### KDM5A/B suppress KSHV lytic reactivation from latency

In light of our early findings of KDM5A/B’s role in silencing HIV proviruses, we were curious whether KDM5A/B impact the lytic/latent switch of KSHV as well. TREx.BCBL-1.RTA cells were electroporated with siRNAs targeting KDM5A, KDM5B, or KDM5C, followed by Dox treatment. KDM5A/B were efficiently depleted (**Fig 2A, 2B**), which significantly promoted the Dox-induced KSHV lytic reactivation (**Fig 2C-F**). However, knockdown of KDM5C showed no such effects (**Fig S2A**, **S2B**). We also confirmed these findings in the iSLK.BAC16 cells, a human renal cell carcinoma cell line with the epithelial cell origin (SLK) that harbors a bacterial artificial chromosome clone (BAC16) of the recombinant KSHV genome while also carrying an integrated cassette for Dox induced RTA expression. Similarly, KDM5A/B depletion by siRNAs dramatically promoted the Dox-induced KSHV lytic reactivation in iSLK.BAC16 cells (**Fig 2G-L**). Such phenotypes were also observed in another iSLK cell model of KSHV, the iSLK.r219 cells harboring the KSHV r219 strain that expresses RFP from KSHV lytic PAN promoter as well as GFP constitutively from the EF1a promoter. KDM5A/B depletion by either siRNAs or shRNAs remarkably increased the RFP florescence intensity in iSLK.r219 cells (**Fig 2M, 2N, S2C, S2D**). Thus, we demonstrated that KDM5A/B but not KDM5C play a role in blocking KSHV lytic reactivation from latency.

**Figure 2.**
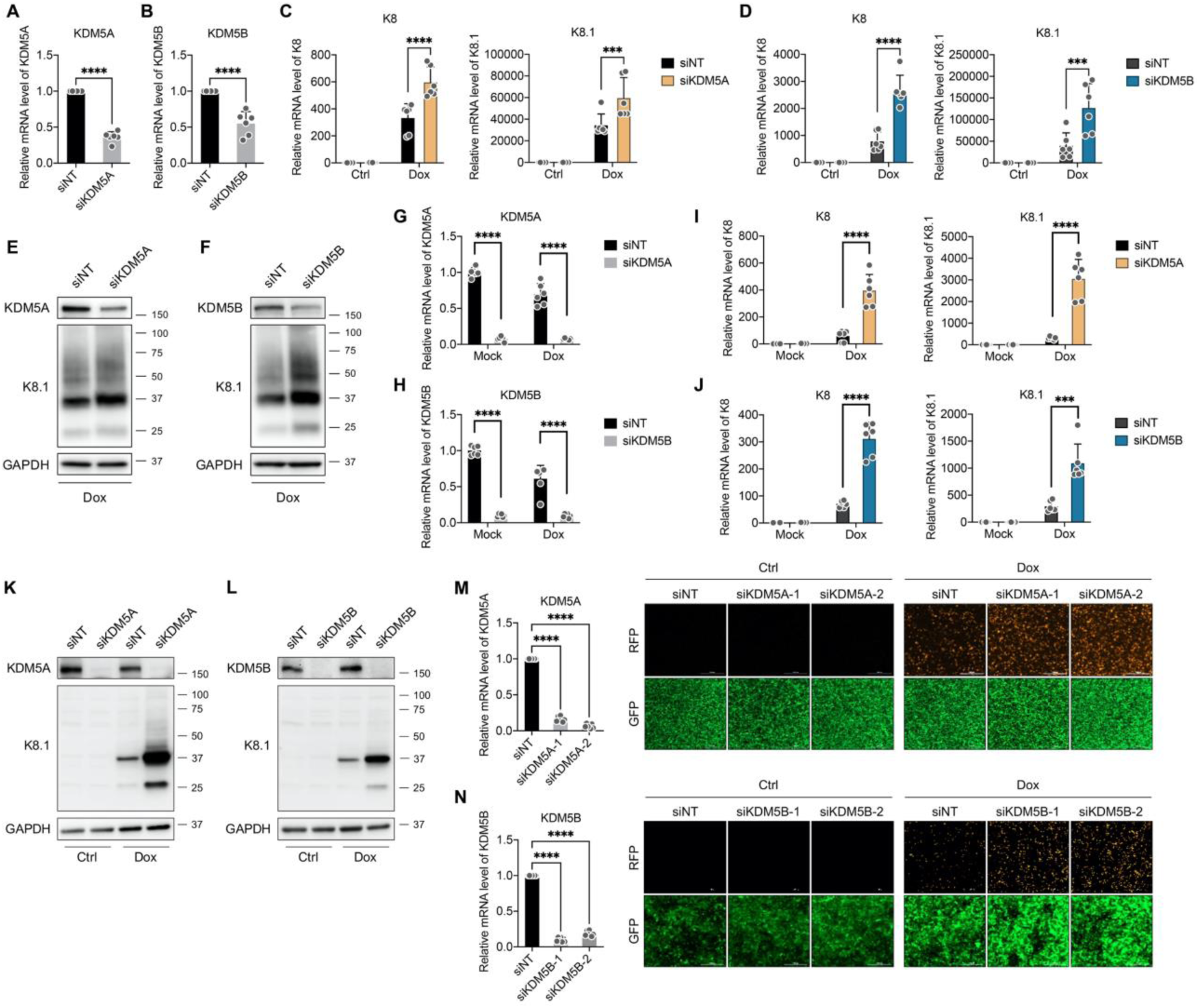
KDM5A/B suppress KSHV lytic reactivation from latency. (A, B) TREx.BCBL-1.RTA cells were transiently transfected with siRNAs targeting KDM5A (A) or KDM5B (B) through electroporation. At 3 days post of transfection, cells were collected and analyzed by RT-qPCR to quantify KDM5A/B knockdown. (C, D) TREx.BCBL-1.RTA cells with depletion of KDM5A (C) or KDM5B (D) were treated with Dox for 2 days, followed by RT-qPCR to quantify KSHV lytic genes K8 and K8.1. (E, F) KDM5A/B and KSHV K8.1 proteins in cell samples (C, D) with Dox treatment were measured by protein immunoblotting. (G, H) iSLK.BAC16 cells were transiently transfected with siRNAs targeting KDM5A (G) or KDM5B (H) for 3 days, followed by Dox treatment for 2 days. KDM5A/B knockdown were measured by RT-qPCR. (I, J) iSLK.BAC16 cells with depletion of KDM5A (I) or KDM5B (J) as well as Dox treatment was analyzed by RT-qPCR to quantify KSHV lytic genes K8 and K8.1. (K, L) KDM5A/B and KSHV K8.1 proteins in cell samples in cell samples (I, J) were measured by protein immunoblotting. (M, N) iSLK.r219 cells were transiently transfected with siRNAs targeting KDM5A (M) or KDM5B (N) for 3 days, followed by Dox treatment for 2 days. KDM5A/B knockdown were measured by RT-qPCR. KSHV lytic reactivation was visualized by fluorescence microscopy (RFP *vs* GFP signal). Results were calculated from three independent experiments and presented as Mean ± SD (*** *P* < 0.001, **** *P* < 0.0001; unpaired, two-tailed Student’s *t*-test for A; one-way ANOVA for I, J, two-way ANOVA for B, C, E, F, G).

### KDM5A/B associate with KSHV LANA and modulate its functions

LANA is a KSHV-encoded oncoprotein critical to maintain viral latency as well as drive viral tumorigenesis(21). A report showed that chromatin association of KSHV LANA overlaps significantly with that of H3K4me3(22), the major substrate of KDM5A/B. We re-analyzed the public-domain chromatin immunoprecipitation followed by sequencing (ChIP-seq) data generated in BC3 cells (GSE202670). We compared the ChIP-seq peaks of H3K4me3 with host (DNA Polymerase I alpha) and KSHV viral proteins (LANA, RTA) extracted from the raw datasets. Average profile plot of ChIP-seq peaks showed the similar patterns only between LANA and H3K4me3 (**Fig 3A**) as opposed to RTA and DNA Polymerase I alpha. We further focused on their binding peaks particularly at loci of immune-responsive genes (IRGs). Profile plots and heatmaps of IRG peaks showed the highly concordant patterns only between LANA and H3K4me3 (**Fig 3B**). The peak overlap enrichment analysis between LANA and H3K4me3 at IRGs was significant with adjusted *P*-value (*P* = 0.0192). These data analyses indicated a cooperative association between LANA and KDM5A/B-targeted H3K4me3 mark to regulate viral and host gene expression, including IRGs. To prove this, we first confirmed that KSHV LANA protein interacts with endogenous KDM5A/B proteins in BCBL-1 cells by protein immunoprecipitations (**Fig 3C**). Such protein interactions were also verified in iSLK.BAC16 cells by both protein immunoprecipitations (**Fig 3D**) and protein colocalization (**Fig 3E, 3F**). KDM5A/B remained to bind with LANA despite that their protein level significantly decreased upon KSHV lytic reactivation in Dox-treated TREx.BCBL-1.RTA (**Fig S3A**) and iSLK.BAC16 (**Fig S3B, S3C**) cells.

**Figure 3.**
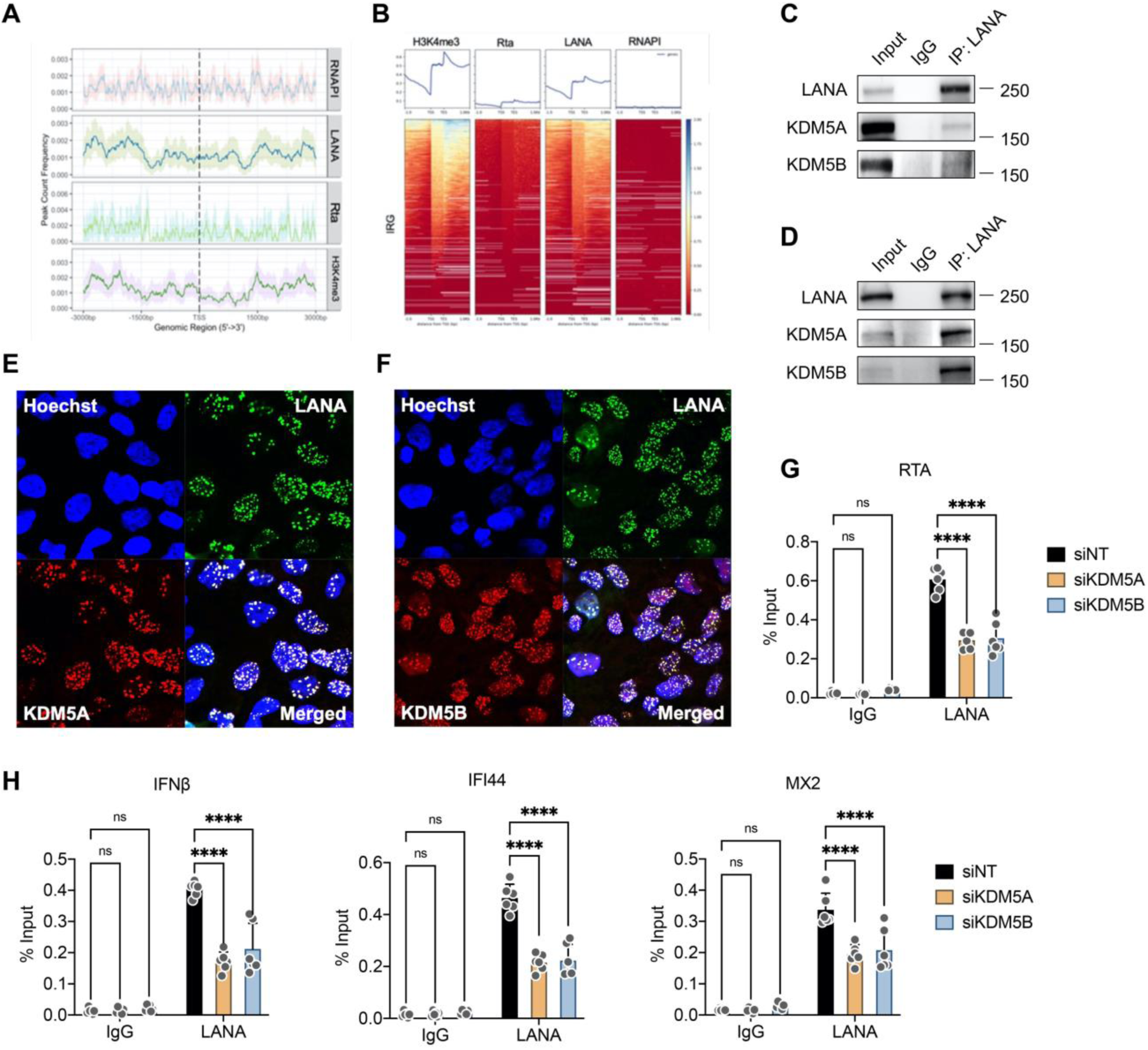
KDM5A/B associate with KSHV LANA and modulate its functions. (A, B) Average profile plot (A) and heatmap (B) showing the ChIP-seq enrichment pattern of RNAP1, LANA (KSHV), RTA (KSHV), and H3K4me3 at promoter regions of immune-responsive genes (IRGs) were presented, which were generated from re-analysis of the public ChIP-seq dataset (GSE202670). (C, D) BCBL-1 (C) or iSLK.BAC16 (D) cells were collected for protein IP of KSHV LANA, followed by protein immunoblotting to quantify the co-IPed KDM5A/B protein. (E, F) iSLK.BAC16 cells were subjected to protein immunofluorescence followed by confocal microscopy imaging to visualize the co-localization of KSHV LANA protein with KDM5A (E) or KDM5B (F). (G, H) iSLK.BAC16 cells were transiently transfected with siRNAs targeting KDM5A/B for 3 days. Cells were collected for chromatin immunoprecipitation (ChIP) of KSHV LANA protein using its antibody or the control rat IgG, followed by qPCR to quantify the binding of LANA at the promoter regions of KSHV RTA (G) or host IRGs (H). Results were calculated from three independent experiments and presented as Mean ± SD (**** *P* < 0.0001; two-way ANOVA).

We next determined whether KDM5A/B interact with LANA to modulate its functions. Depletion of KDM5A/B by their siRNAs only modestly decreased the protein expression of a LANA cDNA transfected in the HEK293T (**Fig S3D**). LANA has been reported to promote KSHV latency through binding to viral lytic promoter that controls RTA expression and suppressing its activities(23–25). We showed that the chromatin association of LANA at RTA promotor significantly decreased in iSLK.BAC16 cells due to KDM5A/B depletion by siRNAs (**Fig 3G**). Furthermore, LANA has also been reported to directly suppress the expression of certain host antiviral genes to favor KSHV viral persistent infection(26,27). It has been implicated that LANA directly associates at the promoter of immune genes to regulate their expression(28), which aligned with our data analysis of LANA ChIP-seq data as well (**Fig 3B**). We demonstrated that KDM5A/B depletion by siRNAs indeed caused a remarkable reduction of LANA’s chromatin association at the promotor of several antiviral immune genes (**(Fig 3H**). Overall, we identified that KDM5A/B associate with KSHV LANA, which showed no obvious impact on LANA protein expression but determined LANA’s chromatin association and its functions to promote KSHV viral latency.

### KDM5 inhibition induces KSHV lytic reactivation from latency

As our own data showed that KDM5A/B prevent latent KSHV from lytic reactivation likely through interaction with LANA to modulate its functions, we speculated that their inhibition would promote KSHV lytic reactivation in KSHV-infected tumor cells and induce the viral oncolysis effect. Recently, a KDM5-specific inhibitor JQKD82 has been developed, which is a prodrug of KDM5-C49(29) and has demonstrated the antitumor potency against multiple myeloma(30). JQKD82 treatment significantly enhanced Dox-induced KSHV lytic reactivation in TREx.BCBL-1.RTA cells by quantifying viral lytic gene expression at the doses with no obvious cytotoxicity (**Fig 4A-C**). Such findings were confirmed by using another KDM5-specific inhibitor CPI-455 that exerted the similar effects on Dox-induced KSHV lytic reactivation in TREx.BCBL-1.RTA cells (**Fig S4A, S4B**). We further tested JQKD82 using KSHV-positive PEL cell lines that harbor latent KSHV, including BCBL-1 and BC-3. JQKD82 alone was competent to potently induce KSHV lytic reactivation in these cells at different doses (**Fig 4D, 4E, S4C, S4D**). Regardless of cell types, JQKD82 still remarkedly promoted Dox-induced KSHV lytic reactivation in iSLK.BAC16 cells (**Fig 4F, 4G**). On the contrary, JQKD82 treatment only slightly decreased the protein expression of a LANA cDNA transfected in HEK293T cells while increasing the total level of H3K4me3 (**Fig S4E**). Therefore, our results illustrated the potential to use KDM5 inhibitors for reversal of latent KSHV that would further lead to viral oncolysis for KSHV-positive B-cell lymphomas.

**Figure 4.**
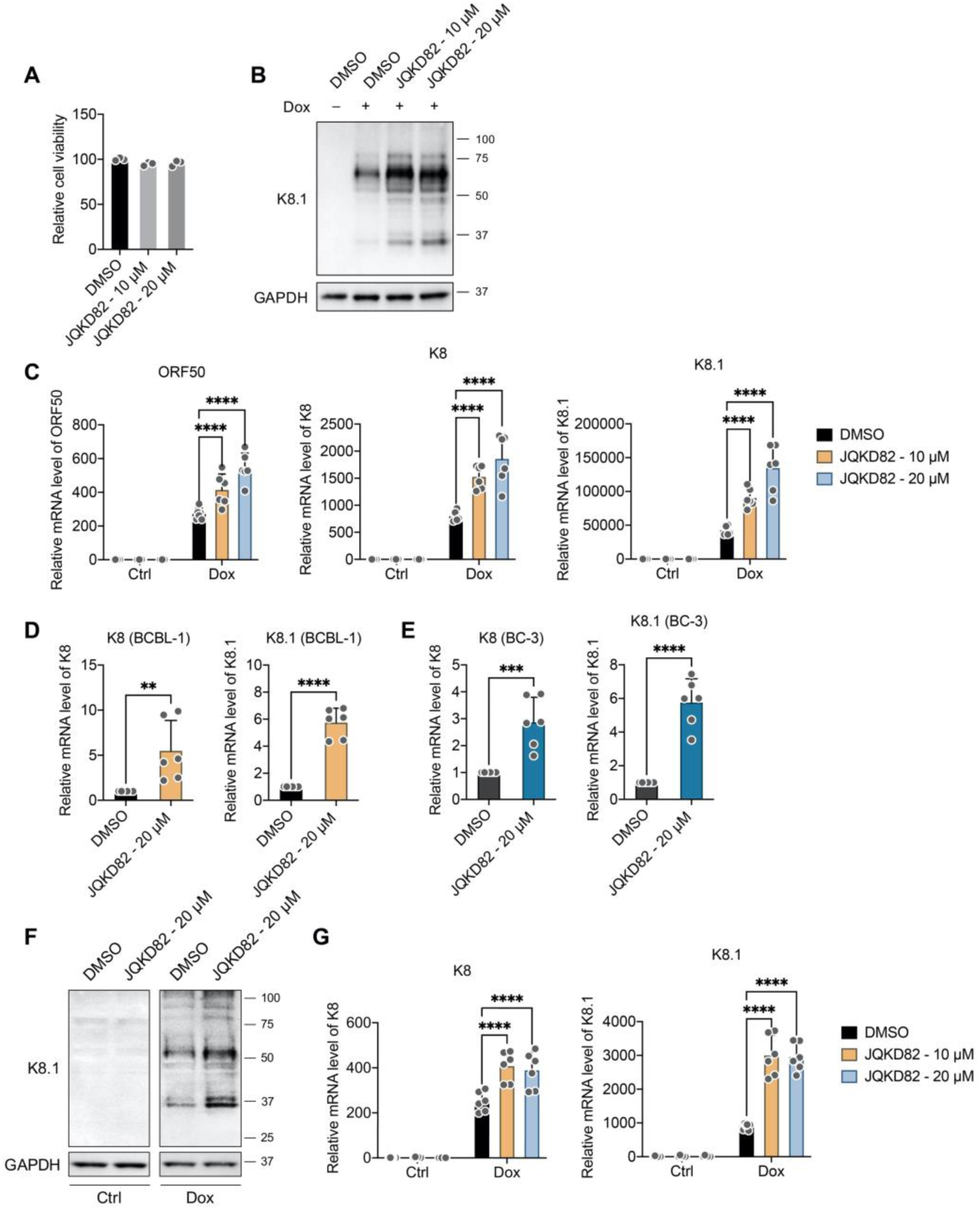
KDM5 inhibition induces KSHV lytic reactivation from latency. (A) TREx.BCBL-1.RTA cells treated with JQKD82 (48 h) was analyzed by the ATP luminescent assay to measure drug’s cytotoxicity. (B) TREx.BCBL-1.RTA cells were treated with JQKD82 for 48 h and then Dox for another 48 h. Cells were collected for protein immunoblotting of KSHV lytic K8.1 protein. (C) mRNA level of KSHV lytic viral genes (ORF50/RTA, K8, K8.1) in cell samples (B) was measured by RT-qPCR. (D, E) BCBL-1 (D) or BC-3 cells (E) were treated with JQKD82 for 96 h and analyzed by RT-qPCR to quantify KSHV lytic genes K8 and K8.1. (F) iSLK.BAC16 cells were treated with JQKD82 for 48 h and then Dox for another 48 h. Cells were collected for protein immunoblotting to quantify KSHV K8.1 protein. (G) iSLK.BAC16 cells treated with JQKD82 in the absence or presence of Dox were collected and analyzed by RT-qPCR to quantify KSHV lytic genes K8 and K8.1. Results were calculated from three independent experiments and presented as Mean ± SD (** *P* < 0.01, *** *P* < 0.001, **** *P* < 0.0001; unpaired, two-tailed Student’s *t*-test for D, E; two-way ANOVA for C, G).

### KDM5 inhibition induces antiviral/antitumor innate immune responses

During KSHV lytic reactivation from latency, host innate immune responses are also activated, which is expected to favor the tumor cell killing by generating the additional oncolytic effects(9). We thus performed the transcriptome profiling analysis for both KSHV-infected BCBL-1 *vs* KSHV-negative BJAB cells with or without JQKD82 treatment by RNA-seq. Although JQKD82 induced the expression of several well-defined antiviral/antitumor IRGs, including IRF6, IRF7, and OAS1, in both BCBL-1 and BJAB cells (**Fig 5A, 5B**), such immune activation was much stronger in BCBL-1 cells indicated by the pathway analysis showing that several immune-related pathways are enriched in JQKD82-treated BCBL-1 cells (**Fig 5C**) but not BJAB (**Fig S5A**).

**Figure 5.**
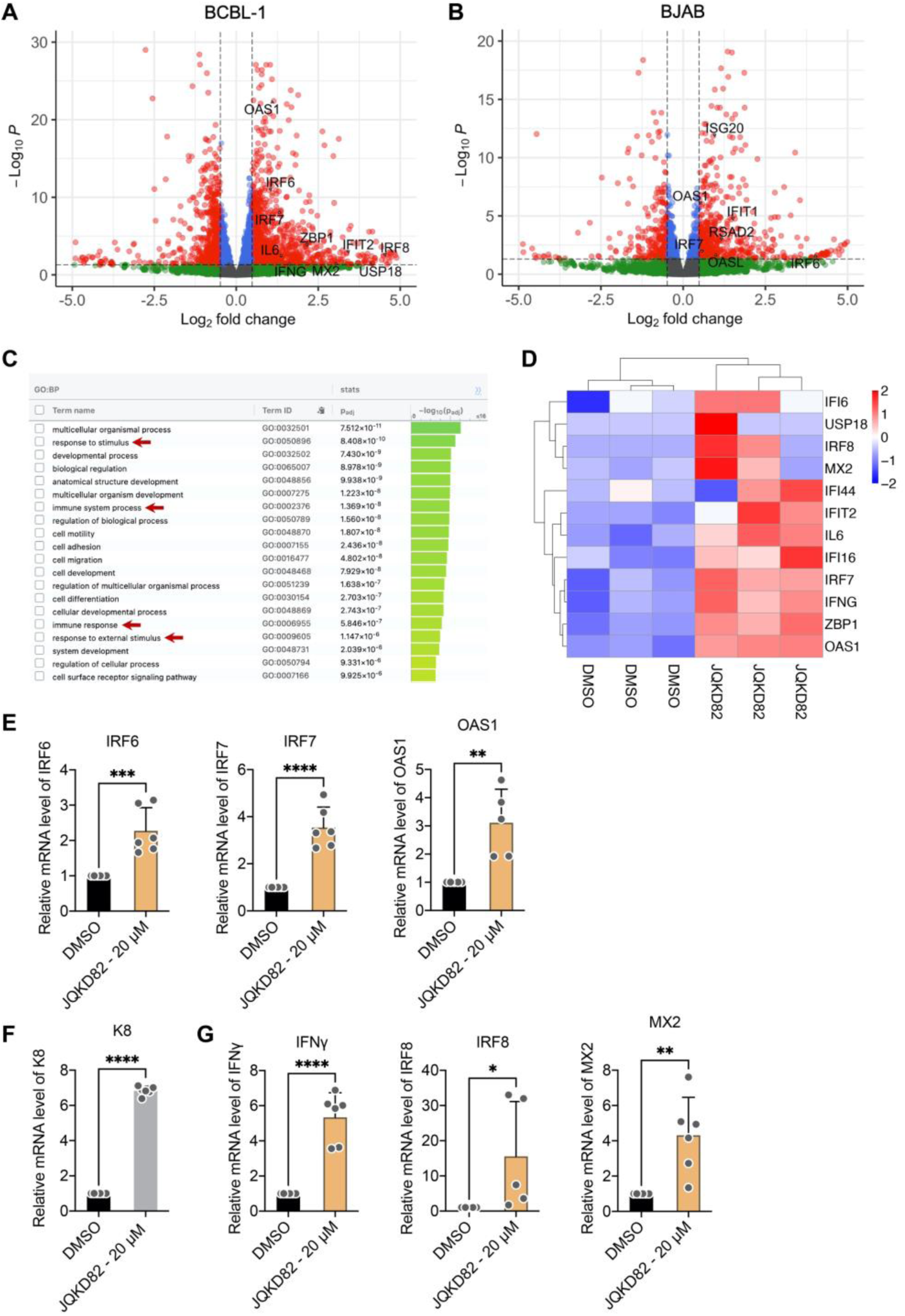
KDM5 inhibition induces antiviral/antitumor innate immune responses. (A, B) BCBL-1 (A) and BJAB (B) cells were treated with JQKD82 (20 µM) for 4 days, followed by RNA-seq analysis. A volcano plot was used to profile the differential gene expression induced by JQKD82 treatment. Several upregulated antiviral/antitumor IRGs were highlighted. (C) GO analysis was performed by using g:Profiler to identify the pathway enrichment for JQKD82-treated BCBL-1 cells. Several immune-related pathways were highlighted. (D) Heatmap was plotted for the selected differentially expressed antiviral/antitumor IRGs in JQKD82-treated BCBL-1 cells. (E-G) BCBL-1 cells were treated with JQKD82 (20 µM) for 4 days, followed by RT-qPCR to quantify the expression of IRGs shared with both JQKD82-treated BCBL-1 and BJAB cells (E), KSHV K8 (F), and IRGs only upregulated in JQKD82-treated BCBL-1 cells (G). Results were calculated from three independent experiments and presented as Mean ± SD (* *P* < 0.05, ** *P* < 0.01, *** *P* < 0.001, **** *P* < 0.0001; unpaired, two-tailed Student’s *t*-test).

We then focused on a set of highly upregulated IRGs in JQKD82-treated BCBL-1 cells for experimental validation by RT-qPCR assays (**Fig 5D**). We first quantified the mRNAs of IRGs identified in both JQKD82-treated BCBL-1 and BJAB cells (IRF6, IRF7, OAS1), which have been reported to suppress tumor growth(31–33). JQKD82 preferentially caused the induction of these IRGs in BCBL-1 (**Fig 5E**) compared to BJAB (**Fig S5B**) cells. We further validated additional antiviral/antitumor IRGs, including IFNγ, IRF8, and MX2, which were enriched only in JQKD82-treated BCBL-1 cells identified from RNA-seq analysis. Along with induction of KSHV lytic gene K8 (**Fig 5F**), these additional IRGs were also preferentially induced in JQKD82-treated BCBL-1 (**Fig 5G**) in comparison to BJAB (**Fig S5C**) cells. Therefore, our transcriptome profiling analysis and further experimental validation suggested that the KDM5 inhibitor JQKD82 is capable of inducing antiviral/antitumor innate immune responses in KSHV-negative B-cell lymphomas, but such effects are more striking in KSHV-positive B-cell lymphomas likely due to KSHV lytic reactivation that further triggers the innate immune activation in tumor cells.

### KDM5 inhibition promotes the death of KSHV-positive PEL cells *in vitro*

Since JQKD82 was capable of inducing viral oncolytic effects and antiviral/antitumor immune responses in KSHV-positive B-cell lymphomas, we next evaluated the drug’s potential to treat these tumors. To establish the clinical relevance of KDM5 proteins as targets for anticancer treatment, we first compared their protein expressions between malignant and normal B cells. In contrast to BJAB cells, KDM5 proteins (KDM5A-C) barely expressed in the CD19^+^CD3^−^ primary B cells isolated from three healthy donors (**Fig 6A, S6A, S6B, S6C**). However, KDM5 protein expressions varied across KSHV-negative BL (BJAB, Ramos) and KSHV-positive PEL (BCBL-1, BC-3) cell lines (**Fig 6B, S6D**). We also performed the protein immunofluorescence assays (IFA) of KDM5A/B using the lymph node tissues from the deidentified, two donors of diffuse large B-cell lymphoma (DLBCL) and two healthy donors, which showed that KDM5A/B proteins highly express in tumorous *vs* normal lymph nodes (**Fig 6C, 6D**). These findings implicated that KDM5 proteins may serve as promising antitumor targets for treating various subtypes of Non-Hodgkin B-cell lymphomas (NHL), including KSHV-associated ones.

**Figure 6.**
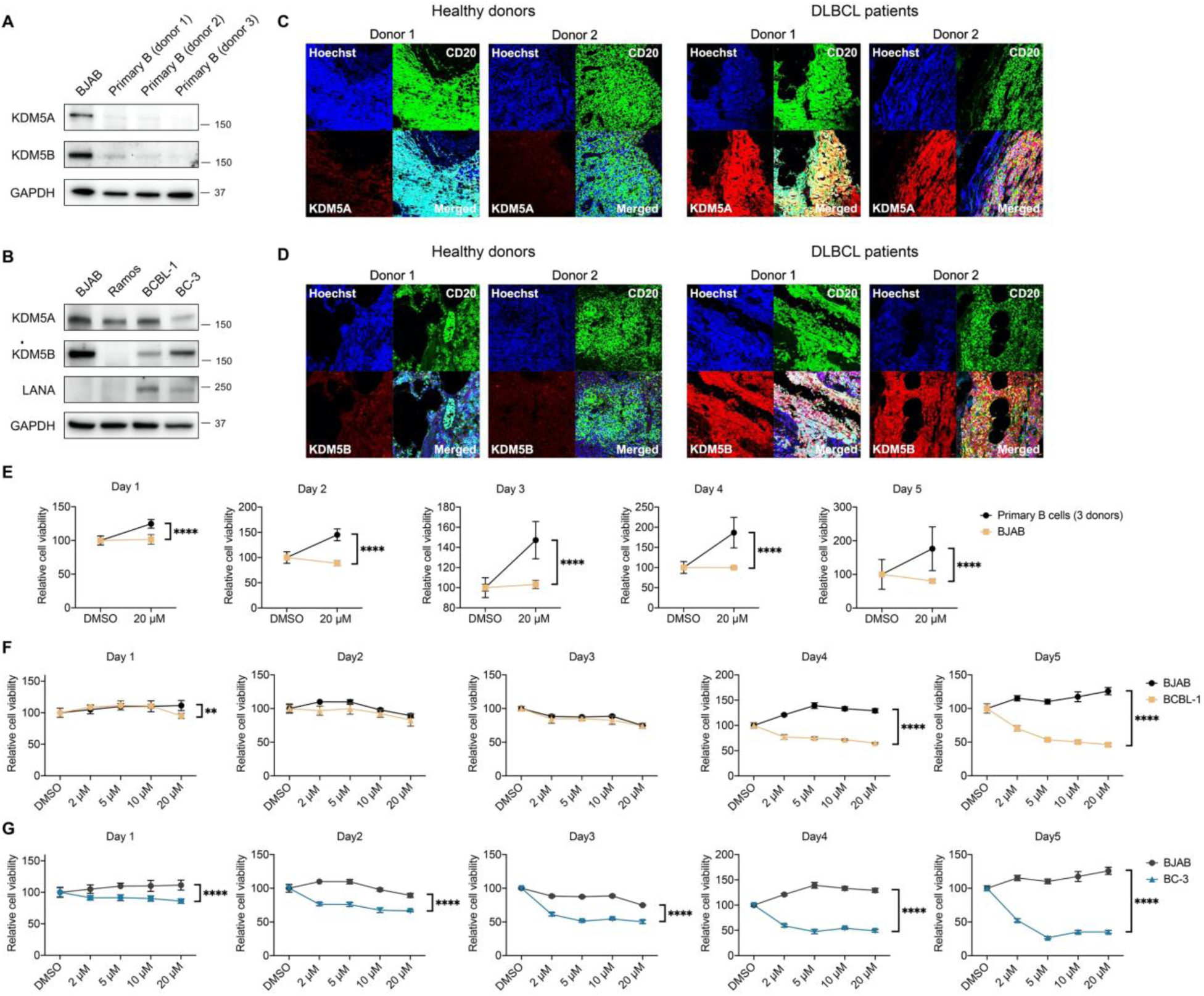
KDM5 inhibition promotes the death of KSHV-positive PEL cells *in vitro*. (A) BJAB and primary B cells (3 heathy donors) were collected and analyzed by protein immunoblotting to quantify KDM5A/B proteins. (B) KSHV-positive (BCBL-1, BC-3) and KSHV-negative (BJAB, Ramos) B-cell lymphoma lines were collected and analyzed by protein immunoblotting to quantify KDM5A/B proteins. (C, D) De-identified lymph node tissue samples (2 healthy donors, 2 diffuse large B-cell lymphoma [DLBCL] patients) were collected and analyzed by protein immunofluorescence to visualize KDM5A (C) or KDM5B (D) protein (Alexa Fluor 647). B-cell marker CD20 was also immuno-stained (Alexa Fluor 488). (E) BJAB and primary B cells (3 heathy donors) were treated with JQKD82 (20 µM) up to 5 days and analyzed by the ATP-based cell viability assay at each day. (F, G) BCBL-1 (F) or BC-3 (G) cells along with BJAB cells were treated with JQKD82 (20 µM) up to 5 days and analyzed by the cell viability assay at each day. Results were calculated from three independent experiments and presented as Mean ± SD (** *P* < 0.01, **** *P* < 0.0001; unpaired, two-tailed Student’s *t*-test).

We then determined JQKD82’s anticancer effects on inducing the cytotoxicity/death of KSHV-positive PEL cells (BCBL-1, BC-3) as well as KSHV-negative BJAB cells using normal B cells as a reference. These cells were cultured up to 5 days with JQKD82 treatment at different doses (5-50 µM). First, our results showed that JQKD82 causes no or only mild cytotoxicity of BJAB cells while even increasing the viability of normal B cells despite of the high doses of JQKD82 (**Fig 6E, S6E**). Second, in contrast to BJAB cells JQKD82 treatment indeed rendered the more obvious cytotoxicity/death of BCBL-1 (**Fig 6F**) and BC-3 (**Fig 6G**) cells across all-tested drug concentrations *in vitro*. Thus, these studies recognized that KDM5 inhibition could be considered as a viable strategy to preferentially treat KSHV-positive B-cell lymphomas.

### KDM5 inhibition blocks the tumor growth of KSHV-positive PEL cells *in vivo*

Given JQKD82 elicited potent antitumor effects on KSHV-positive PEL *in vitro*, we further evaluated this compound on tumor growth *in vivo* using a PEL xenograft mouse model (**Fig 7A**). The NSG mice were transplanted intraperitoneally (i.p.) with the BCBL-1 cells that stably express an integrated cassette of firefly luciferase (BCBL-1-Luc), which allowed the bioluminescence imaging (BLI) to monitor the tumor growth in live animals over the time. The results showed that JQKD82 dramatically inhibits the growth of KSHV-positive PEL in mice during BLI monitoring twice a week up to 28 days post of tumor transplantation in comparison to vehicle control (**Fig 7B, 7C**). The mice were sacrificed, and the tumors and ascites were collected for measurements. JQKD82 treatment substantially decreased the tumor size/volume (**Fig 7D, 7E**), tumor weight (**Fig 7F**), and ascites volume (**Fig 7G**). However, JQKD82 only moderately affected the body weight of PEL-transplanted mice comparing to vehicle control (**Fig S7A**). For the collected tumor samples, we further quantified JQKD82’s drug effects on viral and immune gene expressions by RT-qPCR assays. Consistent with the *in vitro* findings, JQKD82 administration indeed induced KSHV viral gene expressions in tumors (**Fig 7H, S7B**). Likewise, JQKD82 upregulated the expression of anti-tumor IRGs in tumors as well (**Fig 7I, S7C**). Overall, our *in vivo* evaluations clearly demonstrated the therapeutic promise of KDM5 inhibitor JQKD82 to treat KSHV-associated B-cell lymphomas through induction of viral oncolysis.

**Figure 7.**
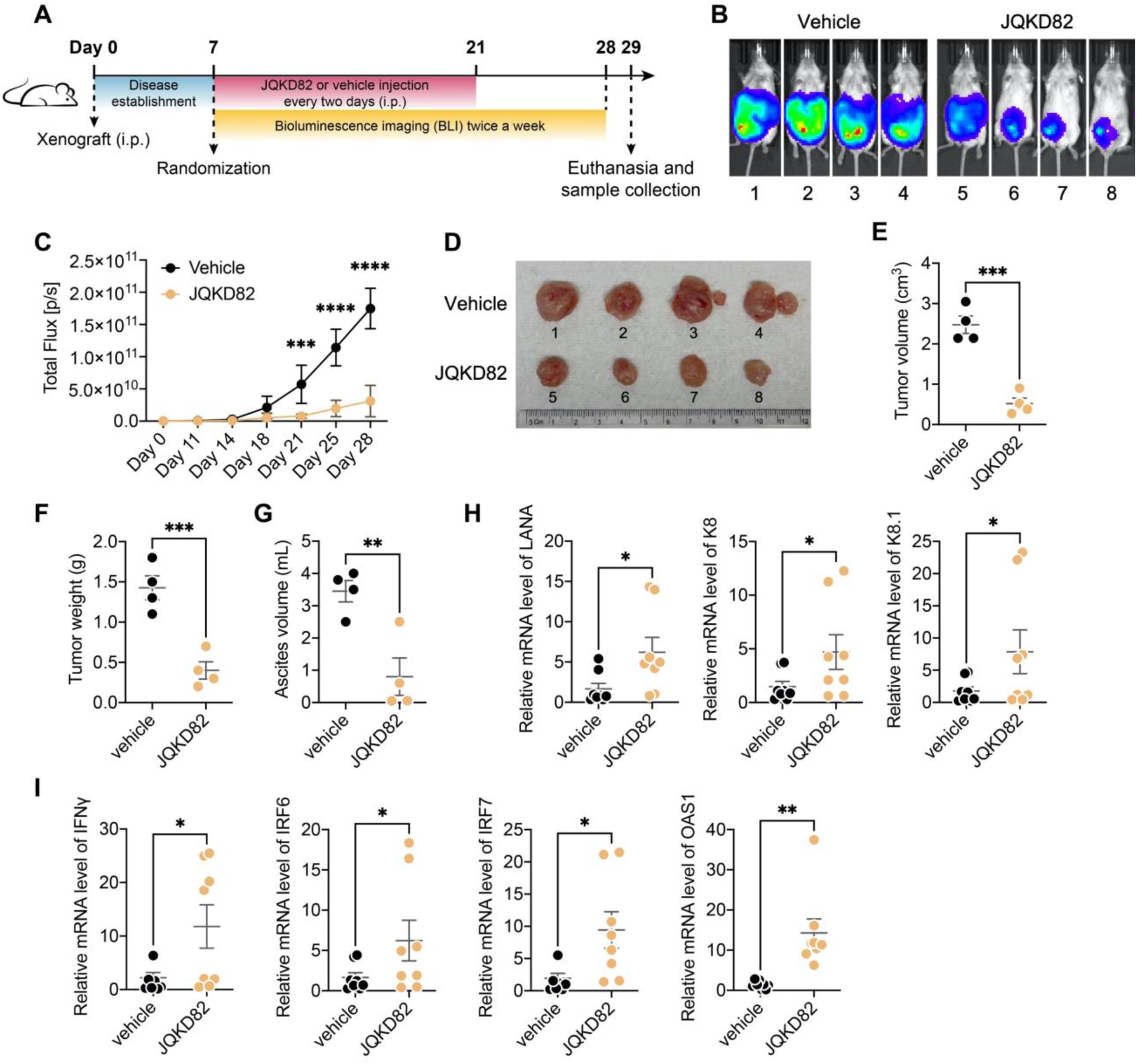
KDM5 inhibition blocks the tumor growth of KSHV-positive PEL cells *in vivo*. (A) A PEL xenograft mouse model was used to evaluate the anti-tumor effect of KDM5 inhibition. Briefly, NSG mice were intraperitoneally (i.p.) transplanted with BCBL-1-Luc cells. At one week post of transplantation, mice were randomized to receive vehicle or JQKD82 that was administrated i.p. at 50 mg/kg every two days for two weeks. Tumor growth was monitored by bioluminescence imaging (BLI) by using the IVIS Spectrum 2 system twice a week for three weeks. (B) Representative images of PEL tumors in either vehicle or JQKD82 mouse group at 28 days post of transplantation were illustrated. (C) Fluc activity (total flux) indicating tumor burden was plotted up to 28 days post of transplantation. (D) Mice were euthanized at Day 29. Tumors were collected from either vehicle or JQKD82 mouse group and photographed for visualization. (E-G) Ascites from tumor sites were also collected from the euthanized mice. Tumor volume (E), tumor weight (F), and ascites volume (G) were measured and compared between vehicle and JQKD82 mouse groups. (H, I) Total RNAs were extracted from harvested tumor samples and analyzed by RT-qPCR to quantify the mRNA level of KSHV viral genes (H) or antitumor IRGs (I). Results were calculated from 4 mice of each condition and presented as either Mean ± SD (for C) or Mean ± SEM (for E, F, G, H, I) (* *P* < 0.05, ** *P* < 0.01, *** *P* < 0.001, **** *P* < 0.0001; unpaired, two-tailed Student’s *t*-test for C; unpaired, one-tailed Student’s *t*-test for E, F, G, H, I).

## Discussion

The viral life cycle of KSHV includes latent and lytic replications as two distinct phases that are delicately controlled by both viral and host factors, assuring KSHV to not only establish persistent infection and escape immune surveillance in the host cells but also generate new progeny viruses once reactivated for viral dissemination and transmission to the new hosts(34). KSHV also encodes an array of viral oncogenes that promote host cell’s transformation and contribute to the development of KSHV-associated malignancies(35). Therefore, elucidation of profound KSHV-host interactions is necessary to advance the understanding of KSHV viral life cycle as well as KSHV-induced tumorigenesis, which would lead to the identification of new gene targets for developing potent and specific anti-KSHV agents. In this study, we provided the novel findings that KDM5A/B proteins play a previously unappreciated role in suppressing KSHV lytic reactivation from latency (**Fig 2**) while KSHV may temporarily downregulate these demethylases via proteosome degradation during its lytic replication to benefit viral propagation (**Fig 1**).

KDM5 family includes four members KDM5A-D, which are the Jumonji-C (JmjC) domain-containing lysine demethylases and catalyze the removal of methyl groups from the epigenetic marks of gene activation H3K4me2/3(36). There is growing evidence showing that KDM5A/B play critical roles in cancer development(15). For example, KDM5A/B are overexpressed in breast cancer(37,38). Loss of KDM5A sensitized the endocrine-resistant luminal breast cancer cells to fulvestrant(39), while loss of KDM5B reduced the proliferation of breast cancer cells *in vivo*(40). Upregulation of KDM5A/B has been observed in multiple types of cancers, such as gastric cancer(41,42), head and neck cancer(43,44), lung cancer(45,46), indicating their strong connections to cancer development. So far, only few reports have linked KDM5A/B proteins to B-cell lymphomas, such as mantle cell lymphoma (MCL) and germinal center (GC) lymphoma(47,48). In contrast, KDM5C/D locate on X and Y chromosomes, respectively, which have been linked to sex-dependent conditions, such as X-linked intellectual disability (XLID), autism, and certain types of sex-specific cancers(49–51). The main B-cell lines we used for this study were mostly generated from BCBL-1 that was derived from an adult male but with an acquired, complete loss of the Y chromosome, while the KSHV-negative BJAB was established from a 5-year-old African girl. Thus, we excluded KDM5D from out studies. Interestingly, we found that although KSHV lytic replication decreases KDM5C protein (**Fig S1B, S1H**), its depletion fails to induce KSHV lytic reactivation (**Fig S2A, S2B**). Since KDM5C may share certain conserved protein domains with KDM5A/B required for KSHV-mediated proteosome degradation, KDM5C is thus also subjected to such dysregulation. However, more likely KDM5A/B play the dominant roles in controlling KSHV viral life cycle.

Our mechanistic investigations identified that KDMA5A/B interaction with LANA may contribute to its functions that promote KSHV viral latency and immune escape to benefit its persistent infection (**Fig 3, S3**). As a pivotal viral factor, KSHV LANA is known to modulate both viral and host gene expression by interacting with cellular proteins, including epigenetic regulators. Early studies have demonstrated that LANA associates with the KSHV viral lytic promoter and negatively regulate its functions to drive the activation of KSHV viral trans-activator RTA that further induces a cascade of KSHV lytic gene expression(23). Indeed, depletion of KDM5A/B significantly decreased LANA chromatin association at the RTA promoter (**Fig 3G**). Furthermore, LANA has also been reported to suppress antiviral/antitumor innate immune responses(26,27). Our data mining provided the supportive evidence that chromatin occupancy of LANA significantly overlaps with that of H3K4me3 at IRG loci (**Fig 3B**). We showed that depletion of KDM5A/B substantially reduces LANA chromatin association at the promoter of IRGs (**Fig 3H**). It is thus plausible that KSHV targets KDM5A/B for proteosome degradation, which would alleviate LANA suppression of RTA activation during viral lytic reactivation. On the other hand, KSHV-mediated KDM5A/B downregulation would also alleviate LANA suppression of antiviral/antitumor IRG activation, resulting in a viral oncolytic impact. Although we ruled out that RTA possesses the functions to induce proteosome degradation of KDM5A/B (**Fig S1J, S1K**), It remains to be identified which viral and/or cellular mechanisms employed by KSHV to do so.

The fundamental understanding of KDM5A/B’s functions to control KSHV viral replications and anti-KSHV innate immune responses would lay the foundation to target them by using small-molecule inhibitors for treating KSHV-associated cancers. KDM5 inhibitors, such as KDM5-C49 or KDM5-C70, have been shown to sensitize endocrine-resistant luminal breast cancer cells to fulvestrant, and the combination of fulvestrant and KDM5 inhibitor substantially reduced tumor volume(39). Although the incidence is rare, KSHV highly associates with certain types of B-cell lymphomas(52), such as PEL, which have the overall lower survival rate than other common types of aggressive B-cell lymphomas(53), leading to short median survival time (only 6-22 months) and poor 5-year survival rate (less than 17%). Although certain treatments extrapolated from other common types of B-cell lymphomas have been applied to PEL and improved its survival rate, there are still no specific therapies developed against these KSHV-associated B-cell lymphomas. Based on our own studies, KDM5A/B would be the ideal targets for treating B-cell lymphomas, as they appeared to preferentially express in malignant *vs* normal B cells (**Fig 6A-D, S6C, S6D**). As a proof of concept, we demonstrated that a recently available KDM5 inhibitor JQKD82 is potent to induce KSHV lytic reactivation and activate antiviral/antitumor innate immune responses in KSHV-infected PEL cells both *in vitro* (**Fig 6**) and *in vivo* (**Fig 7**), facilitating their killing. Such viral oncolytic approaches could be considered for treating other virus-associated cancers. For example, we showed that JQKD82 is capable of inducing lytic reactivation of another human gamma-herpesvirus Epstein-Barr Virus (EBV) and preferentially kills EBV-positive BL cells (BL41-P3HR1) *vs* the parental BL41 (**Fig S7D-F**). However, a limitation is that our NSG mouse model lacks proper immune systems. In future, we will consider using the humanized mouse model with PEL xenograft for further evaluation of KDM5 inhibitor’s impact on antitumor immunity.

## Materials and Methods

### Cells

Cell lines of B-cell lymphomas, including BJAB, Ramos, BL41, BL41-P3HR1, TREx.BJAB.3FLAG.RTA, BCBL-1, BCBL-1-Luc, TREx.BCBL-1.RTA, were maintained in RPMI 1640 medium (Gibco) supplemented with 10% fetal bovine serum (FBS) (Gibco). BC-3 cells were maintained in RPMI 1640 medium supplemented with 20% FBS. iSLK.BAC16, iSLK.r219, HEK293T cells were maintained in Dulbecco’s modified Eagle’s medium (DMEM) (Gibco) supplemented with 10% FBS. Antibiotics were added in cell cultures for TREx.BCBL-1.RTA and TREx.BJAB.3FLAG.RTA (200 μg/mL Hygromycin B), BCBL-1-Luc (50 μg/mL Hygromycin B), iSLK.BAC16 and iSLK.r219 (1.2 mg/mL Hygromycin B, 2 μg/mL puromycin, 250 μg/mL G418) as previously described(54). Telomerase-immortalized human microvascular endothelial (TIME) cells were maintained in Vascular Cell Basal Medium (ATCC, Cat# PCS-100-030) supplemented with Microvascular Endothelial Cell Growth Kit-VEGF (ATCC, Cat# PCS-100-041). Human primary cells, including peripheral blood mononuclear cells (PBMCs) (STEMCELL Technologies) as well as the isolated B cells, were maintained in RPMI 1640 complete medium (RPMI 1640 basic, 1× MEM Non-Essential Amino Acids Solution, 1× Sodium Pyruvate, and 20 mM HEPES) supplemented with human recombinant IL-2 (rIL-2, Roche) at 30 U/mL. Human CD19^+^CD3^−^ primary B cells were isolated from PBMCs using the B cell Isolation Kit II (Miltenyi Biotec, Cat# 130-091-151) according to the manufacturer’s protocol.

### Reagents and antibodies

Doxycycline was purchased from Fisher Scientific (Cat# BP2653-1). DMSO for *in vitro* assays was purchased from Fisher Scientific (Cat# BP231-100). DMSO for *in vivo* applications was purchased from MP Biomedicals (Cat# 2780148). KDM5 inhibitor JQKD82 for *in vitro* experiments was designed, synthesized, and kindly provided by Dr. Jun Qi’s lab at Dana-Farber Cancer Institute(30). JQKD82 for *in vivo* administrations was purchased from Cayman (Cat# 41542). siRNAs targeting KDM5A-C or non-targeting (NT) siRNA were purchased from Invitrogen. Human IgG was purchased from MP Biomedicals (Cat# 55087). MG132 was purchased from Sigma-Aldrich (Cat# 474790). The following antibodies were used in this study: anti-KDM5A (Active Motif, Cat# 91211); anti-KDM5B (Bethyl Laboratories, Cat# A301-813A); ant-KDM5C (Bethyl Laboratories, Cat# A301-034A); anti-KSHV K8.1 (Santa Cruz Biotechnology, Cat# sc-65446); anti-EBV EBNA1 (Santa Cruz Biotechnology, Cat# sc-81581); anti-GAPDH (Santa Cruz Biotechnology, Cat# sc-47724); normal rat IgG (Santa Cruz Biotechnology, Cat# sc-2026); anti-KSHV LANA (Advanced Biotechnologies, Cat# 13-210-100); anti-FLAG M2 (Sigma-Aldrich, Cat# F1804); anti-CD3-FITC (Miltenyi Biotec, Cat# 130-113-690); anti-CD19-PE (Santa Cruz Biotechnology, Cat# 130-113-731); anti-CD20 (BioLegend, Cat# 382802); anti-mouse HRP-linked (Cell Signaling Technology, Cat#7076S) and anti-rabbit HRP-linked (Santa Cruz Biotechnology, Cat# 7074S) antibodies; isotype control IgG including rabbit IgG (Santa Cruz Biotechnology, Cat# 3900S), mouse IgG_1_ (Santa Cruz Biotechnology, Cat# 5415S), mouse IgG_2a_ (Santa Cruz Biotechnology, Cat# 61656S) and mouse IgG_2b_ (Santa Cruz Biotechnology, Cat# 53484S).

### Protein immunoblotting and immunoprecipitation assays

Protein immunoblotting and immunoprecipitation (IP) assays were performed as previously described(55). Briefly, total proteins were extracted from cell lysates by using 1 × radioimmunoprecipitation assay (RIPA) buffer (MilliporeSigma, Cat# 20-188) containing broad-spectrum protease inhibitors (Thermo Scientific, Cat# A32953). Protein concentrations were measured using a BCA assay kit (Thermo Scientific, Cat# 23227), followed by electrophoresis and dry electro-transfer. The PVDF membrane was blocked with nonfat milk, and incubated with primary and HRP-conjugated secondary antibodies, followed by the incubation with ECL substrate for chemiluminescence. To determine protein-protein interactions, cell lysates were incubated with the specific antibodies recognizing the targeted proteins or control IgG, followed by the incubation with protein A/G magnetic beads (Thermo Scientific, Cat# 88803). Beads containing protein immunocomplexes were washed, eluted, and subjected to protein immunoblotting assay. Gel images were acquired by the ProteinSimple FluorChem E Imaging System.

### Protein immunofluorescence assay (IFA)

Primary B cell isolations were evaluated by IFA, followed by flow cytometry analysis as previously described(56). Original PBMCs as well as isolated B cells were incubated with anti-CD3-FITC and anti-CD19-PE antibodies for 30 mins, which were subjected to flow cytometry analysis using BD Accuri C6 Plus. B cell population (CD19^+^/CD3^−^) was analyzed by the FlowJo V10 software. KSHV LANA and KDM5A/B protein interactions were also determined by IFA. iSLK.BAC16 cells were seeded at 1.5 × 10^5^ cells per well on a 24-well culture plate containing circular cover glasses (13 mm) 1 day prior to the induction of KSHV lytic reactivation. Cells were treated with Dox (1 µg/mL) for 2 days. Cells were then fixed with 4% paraformaldehyde at room temperature (RT) for 10 min, followed by permeabilization with 0.2% Triton X-100 at RT for 15 min. The fixed cells were blocked with 1× D-PBS containing 5% FBS for 1 hour at RT and incubated with the primary antibodies (LANA and KDM5A/B) in 1× D-PBS containing 2.5% FBS overnight at 4°C. Anti-rat Alexa Fluor 488 or anti-mouse/rabbit Alexa Fluor 647 conjugated secondary antibodies were then added to cells for another incubation for 1 hour at RT with constant rocking. Nuclei were stained with Hoechst 33342 (Thermo Scientific, Cat# 62249) for 10 min at RT. The cover glasses containing cells were mounted onto a microscope slide with ProLong Glass Antifade Mountant (Invitrogen, Cat# P36980), followed by incubation overnight at 4°C in the dark. Images were acquired using a ZEISS LSM 700 Upright laser scanning confocal microscope with a ZEN imaging software (ZEISS). To quantify KSHV lytic reactivation or *de novo* infection rate, cell samples in culture plates were washed with 1x D-PBS followed by fluorescence imaging using the Cytation 5 multi-mode reader and the Gen5 Image+ software (BioTek).

### Chromatin immunoprecipitation (ChIP)

ChIP assays were performed as previously described(57). In brief, 1% paraformaldehyde (PFA) was used for cell cross-linking. Cells were then re-suspended by CE buffer and centrifuged to pellet nuclei, which were further incubated with SDS lysis buffer and sonicated to shear DNA fragments. 4% of nuclear lysates were used as the Input, and the remaining lysates were incubated with antibodies recognizing the targeted proteins or control IgG, followed by the incubation with pre-blocked protein A/G magnetic beads. The beads were sequentially washed with low-salt buffer, high-salt buffer, LiCl buffer, and TE buffer, and the IPed protein-DNA complexes were isolated by elution buffer. To recover DNA samples, the elutes were treated with 5 M NaCl and incubated at 65 °C for overnight, followed by the treatment of EDTA, Tris-HCl (pH 6.5), and proteinase K. DNA samples were extracted by phenol/chloroform/isoamyl alcohol (25:24:1). DNA pellets were re-suspended in nuclease-free water, which were used for qPCR analysis. The primers used for ChIP-qPCR assays were listed in the supplementary table **(Table S1)**.

### Cell viability assay

ATP-based CellTiter-Glo Luminescent Cell Viability Assay (Promega) was used to measure drug-induced cytotoxicity following the manufacturer’s instructions. Luminescence was measured by the Cytation 5 multimode reader (BioTek).

### KSHV viral propagation and infection

KSHV BAC16 viruses were propagated as previously described with slight modifications(54,56). In brief, iSLK.BAC16 cells were treated with 2 μg/mL doxycycline together with 1 mM sodium butyrate (NaB) for 24 h to reactivate KSHV. Cells were then maintained in fresh medium containing 2 μg/mL doxycycline, and the supernatants containing KSHV BAC16 viruses were harvested every 2 days. Total five batches of harvested supernatants were collected, centrifuged (400 × g) for 10 mins to remove cell debris, and filtered through the 0.45 μm filter. KSHV BAC16 viruses were titrated in HEK293T cells through the serial dilutions. Pre-seeded cells were infected with KSHV BAC16 viruses via spinoculation (2500 rpm) for 2 h at 37 °C.

### Protein immunohistochemistry (IHC) assay

The lymph node specimens from deidentified healthy donors or DLBCL patients were fixed with 4% paraformaldehyde (PFA) at 4°C for overnight, dehydrated through a series of grade ethanol, and incubated with Histo-Clear (National Diagnostics, Cat# 5989-27-5) at room temperature for 2 h prior to paraffin embedding. 7 μm thick sections were then prepared from the paraffin blocks and mounted on the slides. Such paraffin-embedded tissue slides were baked in a high temperature incubator at 65 °C for 30 min and deparaffinized in xylene. The slides were sequentially rehydrated in 100%, 95%, 70%, 50% ethanol and PBS, followed by the antigen unmasking using a pressure cooker with 1 × antigen retrieval buffer (Abcam, Cat# AB93684) for 2 h to complete the cycle and then cool down. Slides were permeabilized with 0.5% Triton X-100 in PBS for 10 min and blocked with 10% normal goat serum (NGS) in 1x PBST for 2 h at room temperature, then incubated with anti-KDM5A, anti-KDM5B, and anti-CD20 antibodies in 5% NGS with 1x PBST at 4°C for overnight. Fluorophore-conjugated secondary antibodies (Alexa Fluor 488 or Alexa Fluor 647) were diluted in 5% NGS with 1x PBST and incubated with the tissue slides for 2 h at room temperature in the dark. These slides were then stained with Hoechst (1:6000 in PBST, Thermo Scientific, Cat# 62249) for 10 min at room temperature, followed by mounting with coverslips using ProLong Glass Antifade Mountant (Invitrogen, Cat# P36982) and curing for overnight. Images were acquired using the ZEISS LSM 700 Upright laser scanning confocal microscope and ZEN imaging software (ZEISS).

### Quantitative reverse transcription PCR (RT-qPCR)

Total RNAs were extracted from cells by using the NucleoSpin RNA isolation kit (MACHEREY-NAGEL, Cat# 740955.250). Eluted RNA samples were reverse transcribed by using the iScript cDNA Synthesis Kit (Bio-Rad, Cat# 1708891) following the manufacturer’s instructions. RT-qPCR assays were performed by using the iTaq Universal SYBR Green Supermix (Bio-Rad, Cat# 1725125) on a Bio-Rad CFX Connect Real-Time PCR System. The primers used for RT-qPCR assays were listed in **Table S1**.

### RNA-seq analysis

BCBL-1 and BJAB cells were treated with JQKD82 (20 μM) or DMSO for 4 days. RNA samples from three independent repeats were extracted by using the NucleoSpin RNA isolation kit following the manufacturer’s manual. RNA samples were submitted to Novogene (Sacramento, CA 95817) for further library preparations. RNA samples that passed quality control were subjected to rRNA removal through polyA selection, followed by the generation of cDNA libraries. Sequencing was carried out on an Illumina NovaSeq X Plus platform with the configuration of 2 × 150 bp (paired end). Differential expression of genes was analyzed by DESeq2(58). R packages, EnhancedVolcano, pheatmap and clusterProfiler programs were used for volcano plotting, heatmap construction and pathway analysis, respectively. The differentially expressed genes (DEGs) in BCBL-1 and BJAB cells due to JQKD82 treatment were listed in the supplementary tables **(Table S2, S3),** respectively. These RNA-seq datasets were deposited in the Gene Expression Omnibus (GEO) with the accession number GSE310839.

### A PEL xenograft mouse model

A PEL xenograft mouse model was employed to determine the antitumor effect of a KDM5 inhibitor JQKD82 on the growth of KHSV-infected PEL cells *in vivo*. Six-week-old female NOD.Cg-*Prkdc^scid^ Il2rg^tm1Wjl^*/SzJ (NSG) mice were acquired from the Ohio State University Comprehensive Cancer Center and transplanted with 5×10^6^ of BCBL-1-Luc cells intraperitoneally (i.p.) per mouse. One week post of tumor xenograft, mice were injected i.p. with D-Luciferin (GoldBio, cat# LUCK-1G) and subjected to the bioluminescence imaging (BLI) for monitoring tumor growth using the IVIS Spectrum 2 In Vivo Imaging System (Revvity). These mice were randomized and grouped for administration with either vehicle control or JQKD82 (50 mg/kg) every two days for two weeks (4 mice per group). The JQKD82 solution was prepared as previously described(30). Briefly, JQKD82 was dissolved in DMSO to prepare the 50 mg/mL solution, which was then further diluted in the 2-Hydroxypropyl)-β-cyclodextrin (Sigma-Aldrich, cat# 332607) with 1:9 ratio (5 mg/mL) for *in vivo* administration. Tumor burden in these mice was monitored by BLI up to three weeks. All mice were euthanized, and tumor samples were collected for further measurements. These animal studies were approved by the Ohio State University Institutional Animal Care and Use Committee (IACUC).

### Statistical analysis

The experimental data were acquired from at least three independent repeats, and analyzed by using either the unpaired, two-tailed/one-tailed Student’s t test or the one-way/two-way analysis of variance (ANOVA). A *P*-value less than 0.05 was considered as statistically significant (\**P* < 0.05, \*\**P* < 0.01, \*\*\**P* < 0.001, \*\*\*\**P* < 0.0001). Results were presented as either Mean ± standard deviation (SD) or Mean ± standard error of the mean (SEM), and graphed by using the GraphPad Prism 9.0 software.

## Acknowledgments

We would like to thank Dr. Renfeng Li (University of Pittsburgh) for providing BJAB cells, Dr. Jae Jung (Cleveland Clinic) for TREx.BCBL-1.RTA cells, Dr. Zsolt Toth (University of Florida) for TREx.BJAB.3FLAG.RTA cells, Dr. Shou-Jiang Gao (University of Pittsburgh) for iSLK.BAC16, iSLK.r219, and TIME cells, Dr. Erle Robertson (University of Pennsylvania) for pA3M-LANA plasmid, and Dr. Robert Baiocchi (Ohio State University) for BL41 and BL41-P3HR1 cells. This study was supported by NIH research grants R01CA260690 to N.S.; R01MH134402, R01DA059538, and R56AI181631 to J.Z.

## Author contributions

N.G.S., J.Z. and D.Z. conceived and designed this study; D.Z. performed the majority of experiments; G.N.F. and S.E. provided experimental supports; Z.W. performed the RNA-seq data analysis; D.Z., N.G.S. and J.Z. analyzed the results; Y.P., J.H., K.A.S.N.S., T.T.L., Y.L., J.O., K.L., J.U.J., J.Q., W.J.Z. contributed materials and/or provided advice for this study; D.Z., N.G.S., J.Z. wrote the manuscript; N.G.S. and J.Z. supervised the entire study.

## Data and materials availability

All data are presented in the main manuscript and/or as the supplementary materials. All other related data, information, and materials can be acquired through the direct requests to authors.

