## Supplementary material for "Inhibition of KDM5A/B promotes antitumor innate immune responses in HHV-8/KSHV-positive B-cell lymphomas": SupplFigures

**Supplementary Figures**

**Figure S1.** (A) TREx.BCBL-1.RTA cells were treated with doxycycline (Dox, 1 μg/mL) for 2 days, followed by RT-qPCR to quantify KSHV lytic gene expression. (B, C) TREx.BCBL-1.RTA cells were treated as (A) and collected for protein immunoblotting to quantify KDM5C and KSHV K8.1 proteins (B) or RT-qPCR to quantify KDM5C mRNA (C). (D, E) BJAB cells were treated as (A) and collected for protein immunoblotting and RT-qPCR to measure KDM5C protein (D) and mRNA € respectively. (F) TIME cells were spinoculated with KSHV BAC16 viruses. At 2dpi, KSHV infection (GFP signal) was visualized by fluorescence microscopy. (G) TIME cells were infected with KSHV as (F) and collected for RT-qPCR to quantify KSHV viral gene expression. (H, I) TIME cells were infected with KSHV as (F) and collected for protein immunoblotting to quantify KDM5C and KSHV K8.1 proteins (H) or RT-qPCR to quantify KDM5C mRNA (I). (J) TREx.BJAB.3FLAG.RTA cells were treated with Dox for 2 days, followed by protein immunoblotting to quantify KDM5A-C proteins. (K) HEK293T cells were transfected with a vector expressing FLAG-tagged RTA or empty vector for 2 days, followed by treatment of MG132 (10 μM) for another 24 h. Cells were collected for protein immunoblotting to quantify KDM5A/B and Flag-RTA proteins. Results were calculated from three independent experiments and presented as Mean ± SD (ns: not significant, ** $P < 0.01$, **** $P < 0.0001$; unpaired, two-tailed Student's *t*-test).

**Figure S2.** (A, B) TREx.BCBL-1.RTA cells were transiently transfected with KDM5C siRNA through electroporation for 3 days, followed by Dox treatment for another 2 days. Cells were collected for RT-qPCR to quantify mRNA level of KDM5C (A) and KSHV viral lytic genes (B). (C) iSLK.r219 cells were stably transduced with KDM5A shRNAs (1, 2) or non-targeting one (shNT), followed by RT-qPCR to confirm KDM5A knockdown. (D) iSLK.r219 cells stably transduced with KDM5A shRNAs or shNT were treated with Dox for 2 days, followed by fluorescence microscopy to visualize KSHV lytic reactivation (GFP *vs* RFP signal). Results were calculated from three independent experiments and presented as Mean ± SD (ns: not significant, * $P < 0.05$, **** $P < 0.0001$; one-way ANOVA for C, two-way ANOVA for A, B).

**Figure S3.** (A) TREx.BCBL-1.RTA cells were treated with or without Dox for 2 days and subjected to protein IP of KSHV LANA. The protein samples were analyzed by protein immunoblotting to quantify LANA and KDM5A/B proteins. (B, C) iSLK.BAC16 cells were treated with Dox for 2 days and subjected to protein immunofluorescence, followed by confocal microscopy imaging to visualize the co-localization of KSHV LANA protein with KDM5A (B) or KDM5B (C). (D) HEK293T cells were transiently transfected with the plasmid pA3M-LANA for 24 h, followed by transfection of the KDM5A or KDM5B siRNA for another 2 days. Cells were collected for protein immunoblotting to quantify LANA and KDM5A/B proteins.

**Figure S4.** (A) TREx.BCBL-1.RTA cells were treated with CPI455 for 48 h. Cytotoxicity was measured by using the ATP luminescent assay. (B) TREx.BCBL-1.RTA cells were treated with CPI455 for 48 h, followed by Dox treatment for another 48 h. Cells were collected for protein immunoblotting to quantify KSHV K8.1 protein. (C, D) BCBL-1 (C) or BC-3 (D) cells were treated with JQKD82 at the high dose (50 µM) for 4 days, followed by RT-qPCR to quantify KSHV lytic gene expression. (E) HEK293T cells were transiently transfected with the plasmid pA3M-LANA for 24 h, followed by treatment of JQKD82 (20 µM) for another 72 h. Cells were collected for protein immunoblotting to quantify LANA and histone markers H3K4me3 and total

H3. Results were calculated from three independent experiments and presented as Mean ± SD (** $P < 0.01$, *** $P < 0.001$, **** $P < 0.0001$; unpaired, two-tailed Student's *t*-test).

**Figure S5.** (A) GO analysis was performed by using g:Profiler to identify the pathway enrichment for JQKD82-treated BJAB cells. (B, C) BJAB cells were treated with JQKD82 (20 μM) for 4 days, followed by RT-qPCR to quantify mRNA level of selected IRGs (B: IRF6, IRF7, OAS1; C: IFNγ, IRF8, MX2). Results were calculated from three independent experiments and presented as Mean ± SD (* $P < 0.05$, ** $P < 0.01$, *** $P < 0.001$; unpaired, two-tailed Student's *t*-test).

**Figure S6.** (A) Gating strategy for quantifying primary B cells isolated from healthy donor's PBMCs by flow cytometry. CD3 and CD19 were immuno-stained to identify human T and B cells respectively. (B) The purity of primary B cells isolated from 3 healthy donors was presented. (C) BJAB and primary B cells (3 heathy donors) were collected and analyzed by protein immunoblotting to quantify KDM5C protein. (D) KSHV-infected (BCBL-1, BC-3) and un-infected (BJAB, Ramos) B-cell lymphoma lines were collected and analyzed by protein immunoblotting to quantify KDM5C protein. (E) BJAB and primary B cells (3 heathy donors) were treated with JQKD82 at the high dose (50 μM) up to 5 days and analyzed by the ATP-based cell viability assay at each day. Results were calculated from three independent experiments and presented as Mean ± SD (**** $P < 0.0001$; unpaired, two-tailed Student's *t*-test).

**Figure S7.** (A) Body weight of the mice from vehicle and JQKD82 groups was monitored up to 28 days post of PEL (BCBL-1-Luc) transplantation. (B, C) Total RNAs were extracted from tumor samples harvested at Day 29 and analyzed by RT-qPCR to quantify the mRNA level of KSHV viral genes (B: ORF50) or antitumor IRGs (C: MX2, IFIT1). (D) EBV-negative (BJAB, BL41) and EBV-positive (BL41-P3HR1) B-cell lymphoma lines were collected and analyzed by protein immunoblotting to quantify KDM5A-C and EBNA1(EBV) proteins. (E) BL41-P3HR1 cells were treated with JQKD82 (10 or 20 μM) for 4 days *in vitro*. Cells were collected and analyzed by RT-qPCR to quantify the mRNA level of EBV lytic genes, including immediate-early (BZLF1, BRLF1), early (BMRF1, BALF5), and late genes (BcLF1, BLLF1). (F) BL41-P3HR1 and BL41 cells were treated with JQKD82 (2-20 μM) or DMSO *in vitro* up to 4 days and analyzed by the cell viability assay at each day. Results were calculated from three independent experiments and presented as either Mean ± SEM (for B, C) or Mean ± SD (for E, F) (* $P < 0.05$, ** $P < 0.01$, **** $P < 0.0001$; unpaired, one-tailed Student's *t*-test for B, C; unpaired, two-tailed Student's *t*-test for E, F).

#### Figure S1

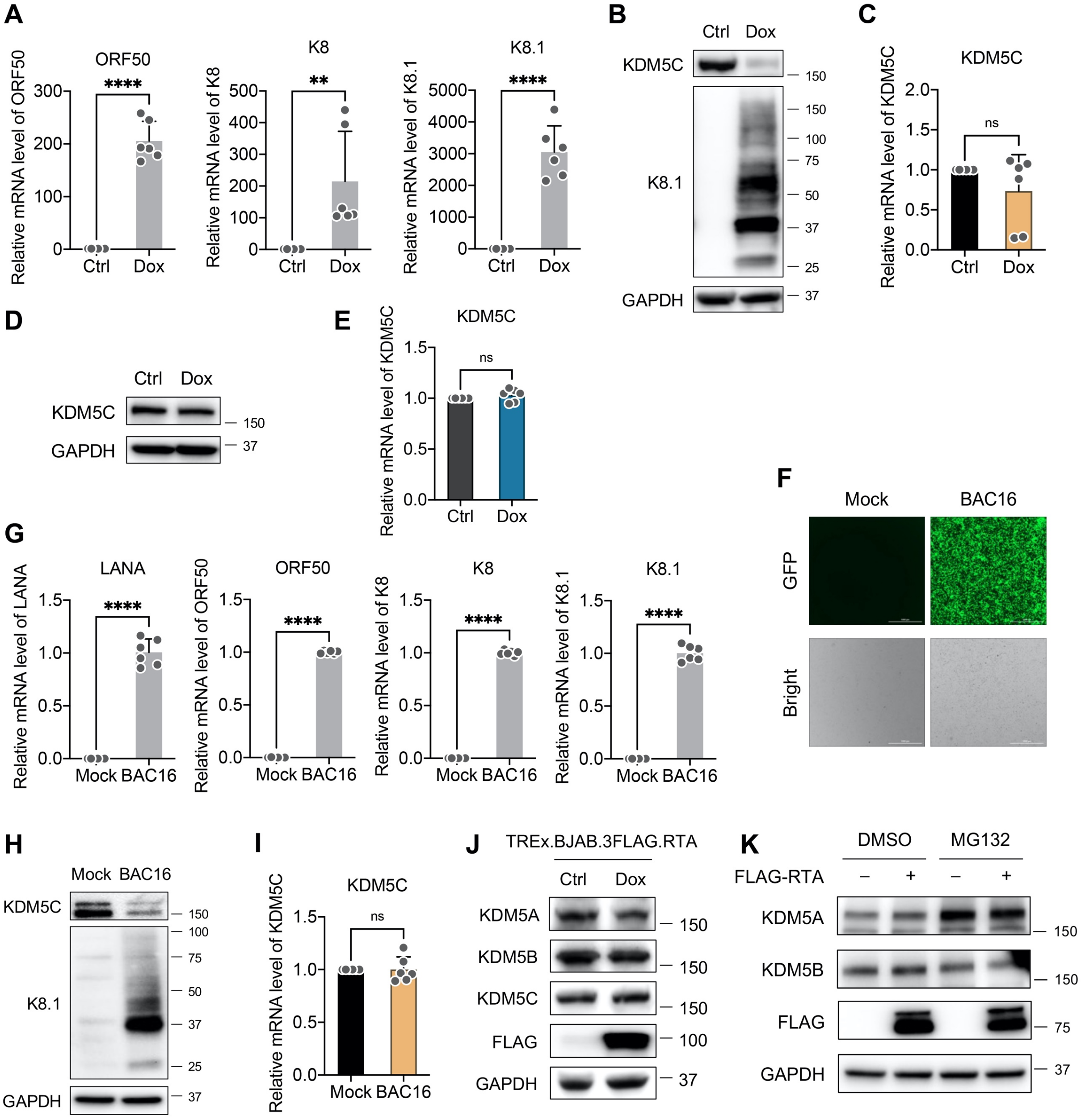
A
ORF50
K8
K8.1
Relative mRNA level of ORF50
Relative mRNA level of K8
Relative mRNA level of K8.1
Ctrl
Dox
B
KDM5C
K8.1
GAPDH
C
KDM5C
Relative mRNA level of KDM5C
ns
D
E
F
Mock
BAC16
GFP
Bright
G
LANA
Relative mRNA level of LANA
H
I
J
TREx.BJAB.3FLAG.RTA
KDM5A
KDM5B
FLAG
K
DMSO
MG132
FLAG-RTA

**Figure S2**

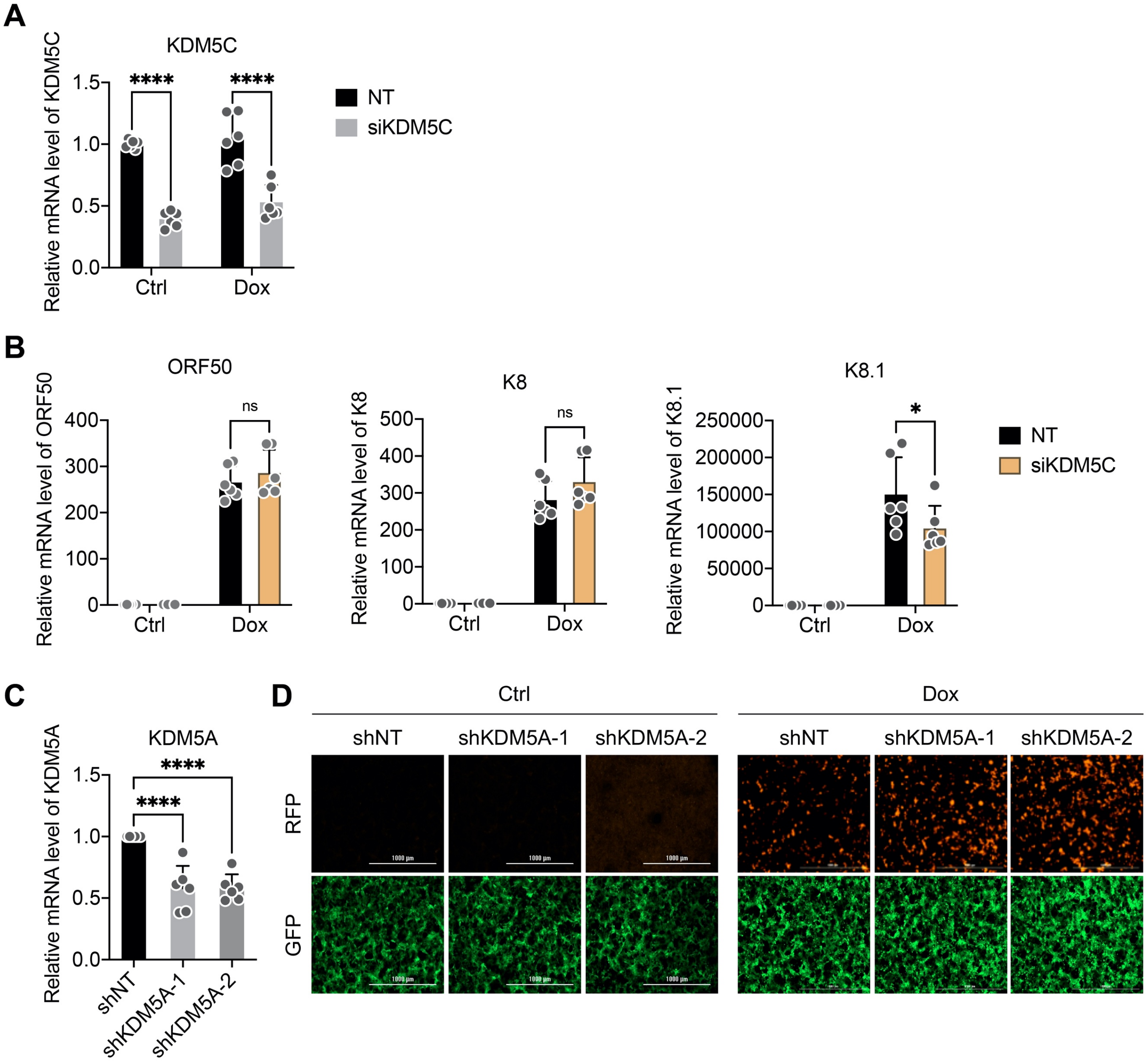
A
KDM5C
Relative mRNA level of KDM5C
1.5
1.0
0.5
0.0
****
****
Ctrl
Dox
NT
siKDM5C
B
ORF50
Relative mRNA level of ORF50
400
300
200
100
0
ns
Ctrl
Dox
K8
Relative mRNA level of K8
500
400
300
200
100
0
ns
Ctrl
Dox
K8.1
Relative mRNA level of K8.1
250000
200000
150000
100000
50000
0
*
Ctrl
Dox
NT
siKDM5C
C
KDM5A
Relative mRNA level of KDM5A
1.5
1.0
0.5
0.0
****
****
shNT
shKDM5A-1
shKDM5A-2
D
Ctrl
Dox
shNT
shKDM5A-1
shKDM5A-2
shNT
shKDM5A-1
shKDM5A-2
RFP
GFP

**Figure S3**

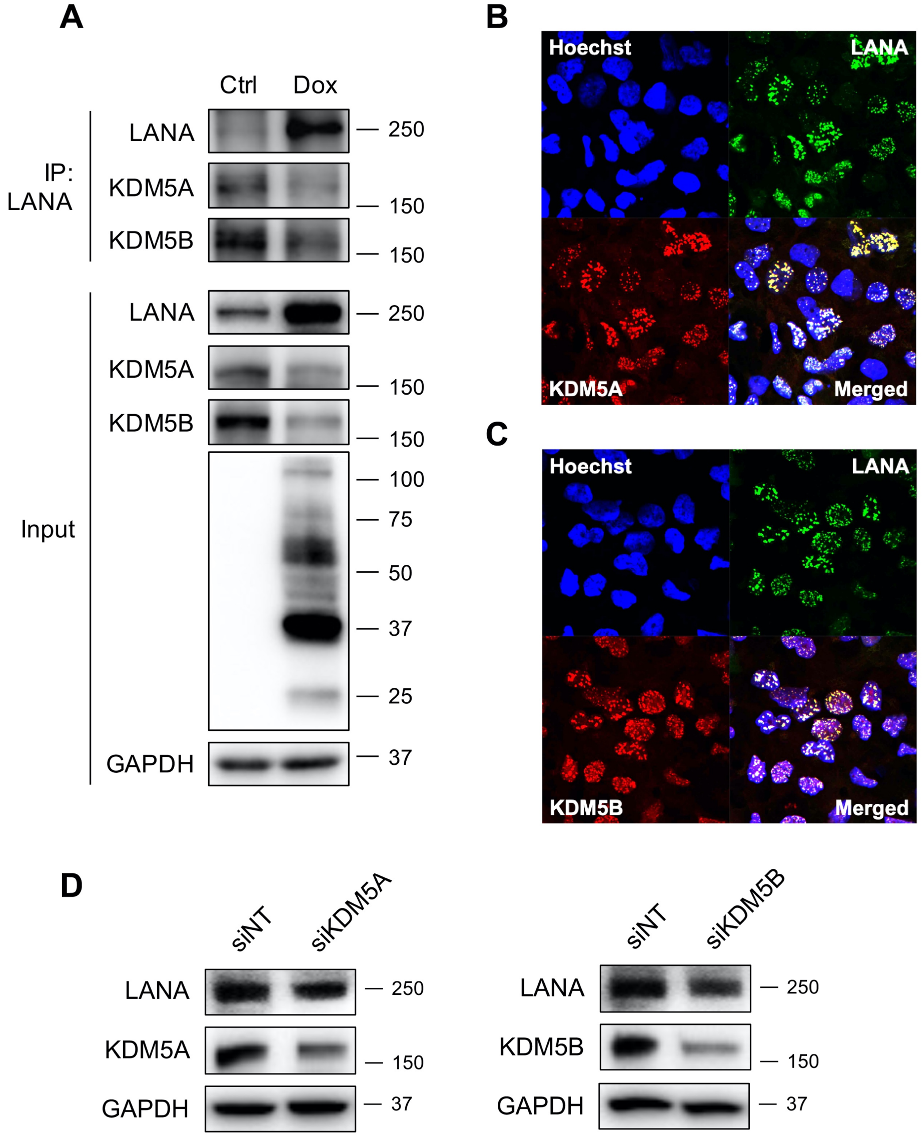
A
Ctrl
Dox
IP: LANA
LANA
KDM5A
KDM5B
Input
LANA
KDM5A
KDM5B
GAPDH
250
150
150
250
150
150
100
75
50
37
25
37
B
Hoechst
LANA
KDM5A
Merged
C
Hoechst
LANA
KDM5B
Merged
D
siNT
siKDM5A
LANA
KDM5A
GAPDH
250
150
37
siNT
siKDM5B
LANA
KDM5B
GAPDH
250
150
37

**Figure S4**

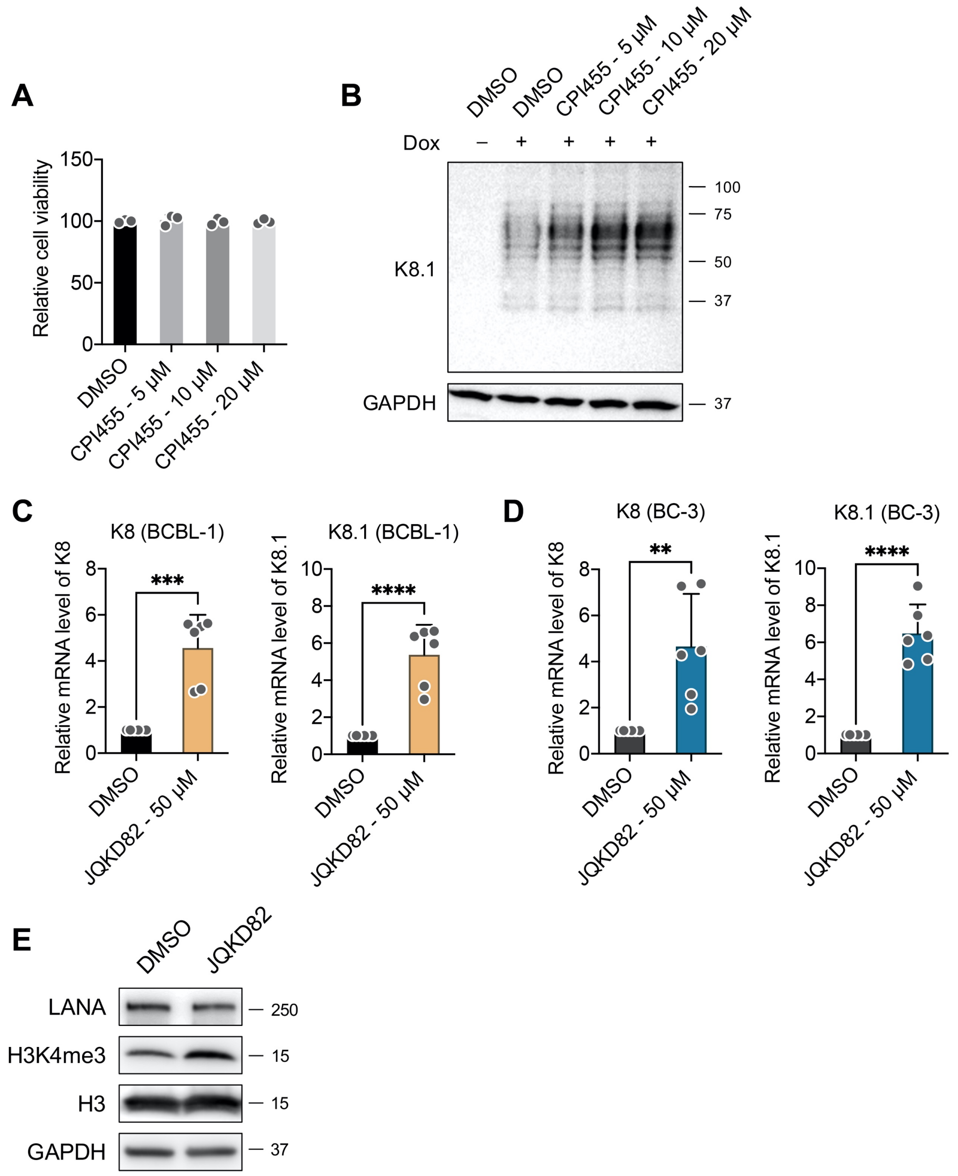
A
Relative cell viability
150
100
50
0
DMSO
CPI455 - 5 µM
CPI455 - 10 µM
CPI455 - 20 µM
B
DMSO
DMSO
CPI455 - 5 µM
CPI455 - 10 µM
CPI455 - 20 µM
Dox
–
+
+
+
+
K8.1
100
75
50
37
GAPDH
37
C
K8 (BCBL-1)
Relative mRNA level of K8
***
K8.1 (BCBL-1)
Relative mRNA level of K8.1
****
DMSO
JQKD82 - 50 µM
D
K8 (BC-3)
Relative mRNA level of K8
**
K8.1 (BC-3)
Relative mRNA level of K8.1
****
DMSO
JQKD82 - 50 µM
E
DMSO
JQKD82
LANA
250
H3K4me3
15
H3
15
GAPDH
37

**Figure S5**

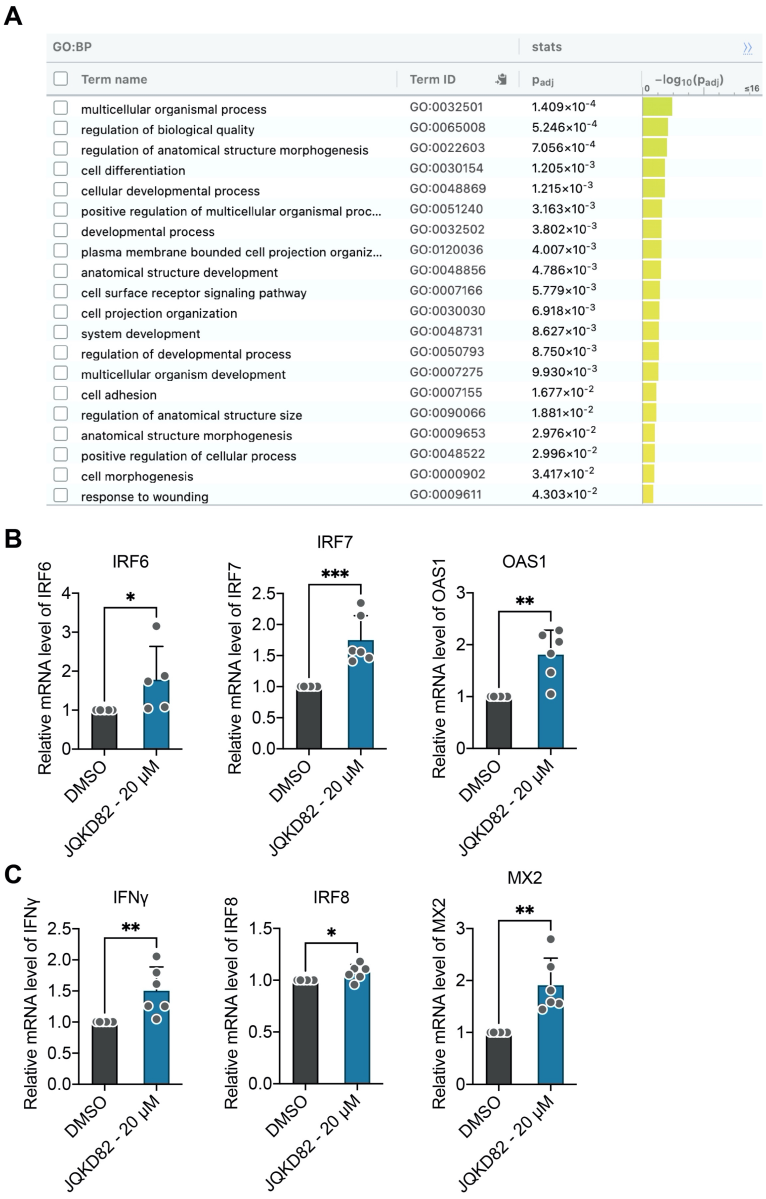

| Term name | Term ID | $p_{adj}$ |
|---|---|---|
| multicellular organismal process | GO:0032501 | $1.409\times10^{-4}$ |
| regulation of biological quality | GO:0065008 | $5.246\times10^{-4}$ |
| regulation of anatomical structure morphogenesis | GO:0022603 | $7.056\times10^{-4}$ |
| cell differentiation | GO:0030154 | $1.205\times10^{-3}$ |
| cellular developmental process | GO:0048869 | $1.215\times10^{-3}$ |
| positive regulation of multicellular organismal proc... | GO:0051240 | $3.163\times10^{-3}$ |
| developmental process | GO:0032502 | $3.802\times10^{-3}$ |
| plasma membrane bounded cell projection organiz... | GO:0120036 | $4.007\times10^{-3}$ |
| anatomical structure development | GO:0048856 | $4.786\times10^{-3}$ |
| cell surface receptor signaling pathway | GO:0007166 | $5.779\times10^{-3}$ |
| cell projection organization | GO:0030030 | $6.918\times10^{-3}$ |
| system development | GO:0048731 | $8.627\times10^{-3}$ |
| regulation of developmental process | GO:0050793 | $8.750\times10^{-3}$ |
| multicellular organism development | GO:0007275 | $9.930\times10^{-3}$ |
| cell adhesion | GO:0007155 | $1.677\times10^{-2}$ |
| regulation of anatomical structure size | GO:0090066 | $1.881\times10^{-2}$ |
| anatomical structure morphogenesis | GO:0009653 | $2.976\times10^{-2}$ |
| positive regulation of cellular process | GO:0048522 | $2.996\times10^{-2}$ |
| cell morphogenesis | GO:0000902 | $3.417\times10^{-2}$ |
| response to wounding | GO:0009611 | $4.303\times10^{-2}$ |

**Figure S6**

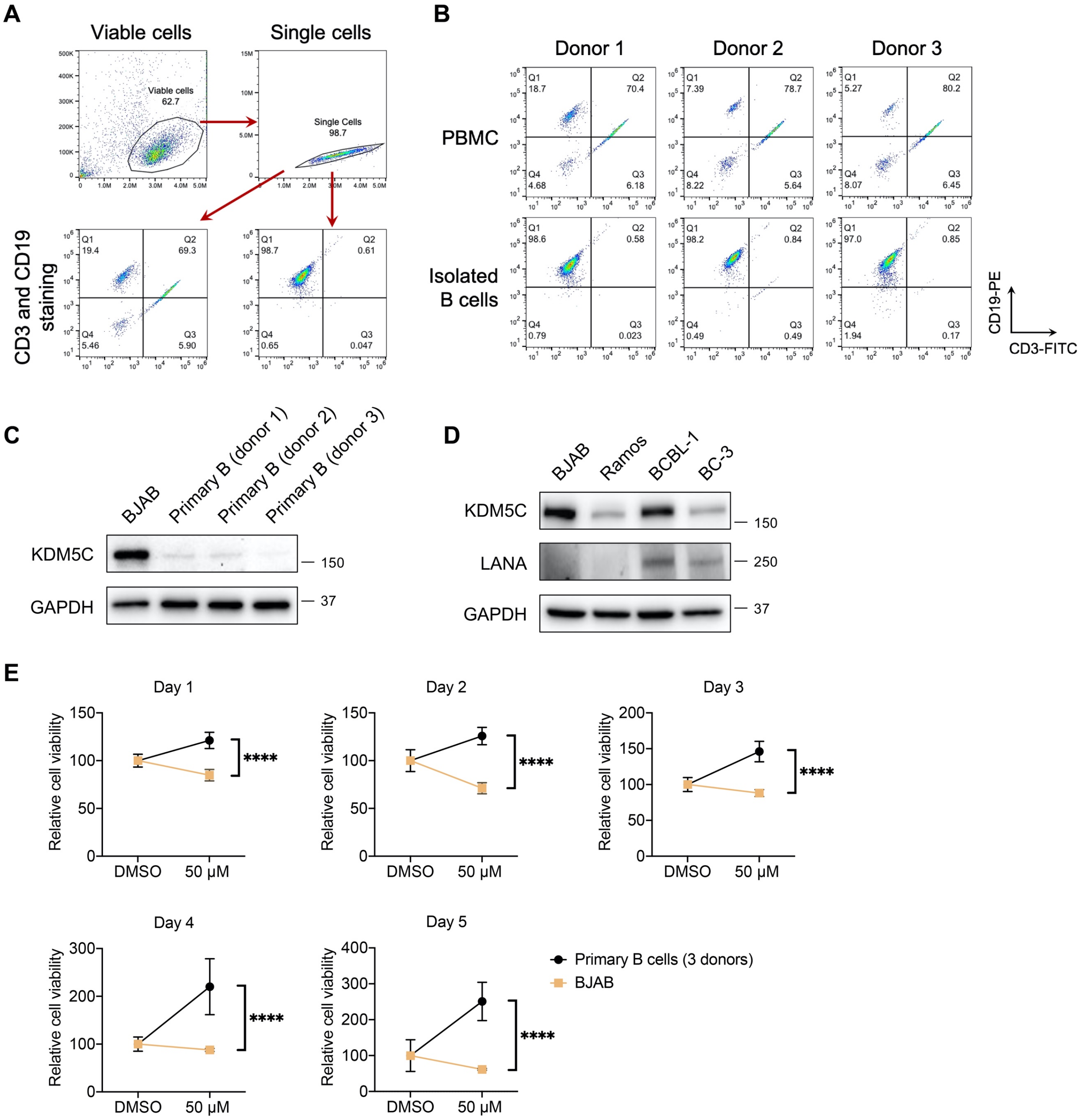
A
Viable cells
Single cells
CD3 and CD19 staining
B
Donor 1
Donor 2
Donor 3
PBMC
Isolated B cells
CD19-PE
CD3-FITC
C
BJAB
Primary B (donor 1)
Primary B (donor 2)
Primary B (donor 3)
KDM5C
GAPDH
D
BJAB
Ramos
BCBL-1
BC-3
KDM5C
LANA
GAPDH
E
Day 1
Day 2
Day 3
Day 4
Day 5
Relative cell viability
DMSO
50 μM
Primary B cells (3 donors)
BJAB

### Figure S7

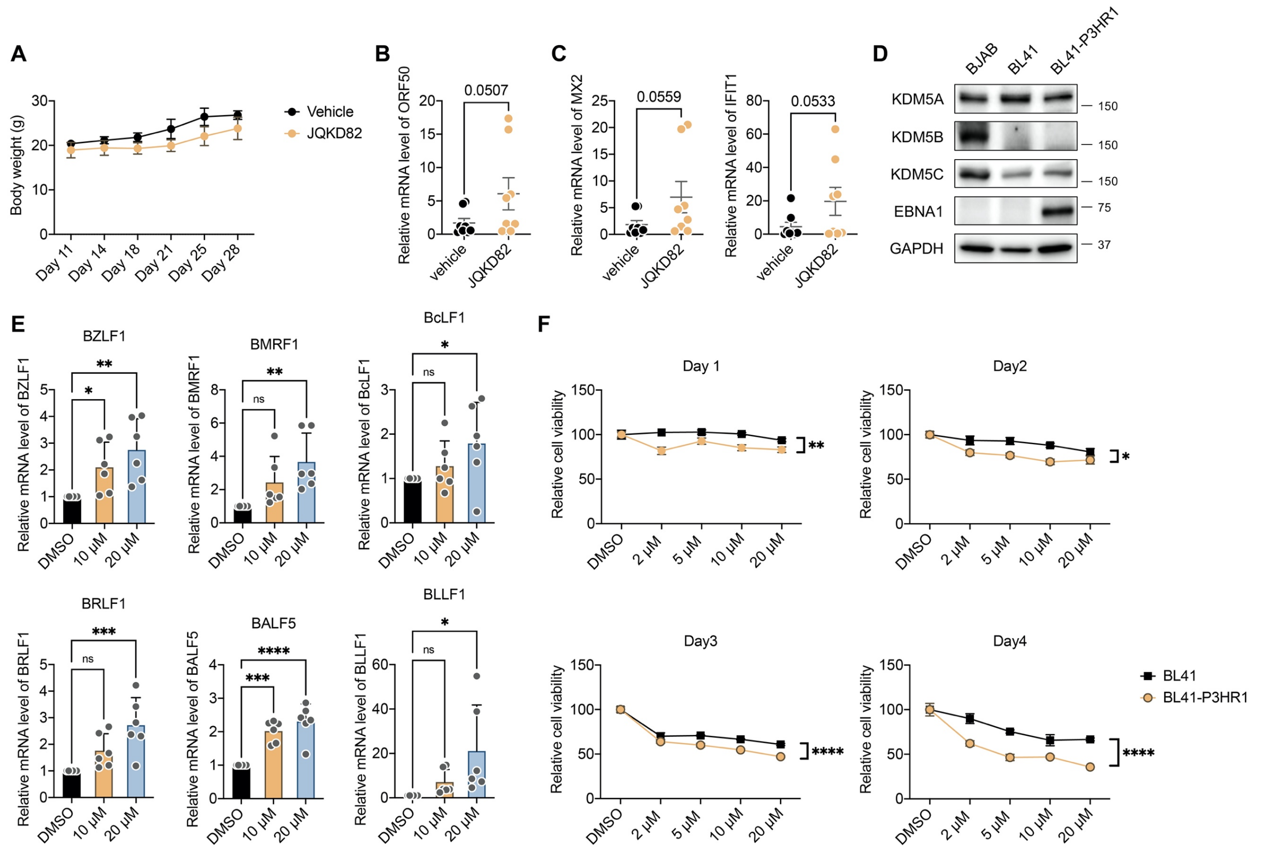
A
B
C
D
E
F
Body weight (g)
Day 11
Day 14
Day 18
Day 21
Day 25
Day 28
Vehicle
JQKD82
Relative mRNA level of ORF50
0.0507
vehicle
JQKD82
Relative mRNA level of MX2
0.0559
Relative mRNA level of IFIT1
0.0533
BJAB
BL41
BL41-P3HR1
KDM5A
KDM5B
KDM5C
EBNA1
GAPDH
150
75
37
BZLF1
BMRF1
BcLF1
BRLF1
BALF5
BLLF1
Relative mRNA level of BZLF1
Relative mRNA level of BMRF1
Relative mRNA level of BcLF1
Relative mRNA level of BRLF1
Relative mRNA level of BALF5
Relative mRNA level of BLLF1
DMSO
10 μM
20 μM
ns
Day 1
Day2
Day3
Day4
Relative cell viability
2 μM
5 μM
BL41
BL41-P3HR1
